# Use of a filter cartridge combined with intra-cartridge bead beating improves detection of microbial DNA from water samples

**DOI:** 10.1101/435305

**Authors:** Masayuki Ushio

## Abstract

Microbial communities play an important role in driving the dynamics of aquatic ecosystems. As difficulties in DNA sequencing faced by microbial ecologists are continuously being reduced, sample collection methods and the choice of DNA extraction protocols are becoming more critical to the outcome of any sequencing study. In the present study, I added a manual, intra-cartridge, bead-beating step in the protocol using a DNeasy® Blood & Tissue kit for DNA extraction from a filter cartridge (Sterivex™ filter cartridge) with-out breaking the cartridge unit (“Beads” method), and compared its performance with those of two other protocols that used the filter cartridge (“NoBeads” method, which was similar to the Beads method but without the bead-beating step, and “PowerSoil” method, which followed the manual of the DNeasy® PowerSoil DNA extraction kit after breaking apart the filter cartridge). Water samples were collected from lake, river, pond and coastal ecosystems in Japan, and DNA was extracted using the three protocols. Then, the V4 region of prokaryotic 16S rRNA genes was amplified. In addition, internal standard DNAs were included in the DNA library preparation process to estimate the number of 16S rRNA gene copies. The DNA library was sequenced using Illumina MiSeq, and sequences were analyzed using the amplicon sequence variant (ASV) approach implemented in the DADA2 pipeline. I found that, 1) the total prokaryotic DNA yields were highest with the Beads method, 2) the number of ASVs (a proxy for species richness) was also highest with the Beads method, 3) overall community compositions were significantly different among the three methods, and 4) the number of method-specific ASVs was highest with the Beads method. These results were generally robust across samples from all aquatic ecosystems examined. In conclusion, the inclusion of a bead-beating step performed inside the filter cartridge increased the DNA yield as well as the number of prokaryotic ASVs detected compared with the other two methods. Performing the bead-beating step inside the filter cartridge causes no dramatic increase in either handling time or processing cost and it can reduce the potential contamination risk from the ambient air and/or other samples. Therefore, this method has the potential to become one of the major choices when one aims to extract aquatic microbial DNAs.

## Introduction

Microbial communities play an important role in driving the dynamics of aquatic ecosystems, including primary production and organic matter decomposition as well as dynamics of macro-organisms (Arrigo 2004, Srivastava et al. 2017). Recent advances in molecular techniques enable the generation of massive amounts of information encoded in microbial environmental DNA at a reasonable cost (e.g., Meyer and Kircher 2010, Jain et al. 2016). For example, Illumina MiSeq/HiSeq/NovaSeq enables generation of tens of millions to billions of sequences, and MiSeq enables the sequencing of a relatively long reads which are preferable when analyzing microbial DNAs (i.e., up to 600 bp long by MiSeq V3 300×2 cycles Sequencing Kit). More recently, Oxford Nanopore Technologies and PacBio provide a way to analyze much longer sequences (e.g., *>* 10 kb by using MinION/GridION/PromethION or RS II/Sequel). All of these technologies enable researchers to detect and identify microbes in nature more easily and accurately than ever before. As a consequence, a wide range of aquatic microbial community studies have utilized molecular techniques (Sharma et al. 2013, Bryant et al. 2016). These trends are also true for microbial communities in non-aquatic ecosystems (Lauber et al. 2009, Caporaso et al. 2011, 2012, Fierer et al. 2011, Ushio et al. 2015).

Sample collection methods and the choice of DNA extraction protocols are becoming more critical to the outcome of any sequencing study in microbial ecology since difficulties of DNA sequencing faced by microbial ecologists are continuously being reduced. In aquatic microbial studies, water sampling and filtration are a first step for studying microbial communities. Use of a filter cartridge (e.g., Sterivex ™ filter cartridge, Merck Millipore) enables on-site filtration in a convenient way, and thus can avoid unnecessary degradation of target microbial DNA during transport (filter cartridges have been used in many studies; e.g., Somerville et al. 1989, Sharma et al. 2013, Bryant et al. 2016, Perrine et al. 2017). In addition, the filter is encapsulated, and thus the risk of contamination can be reduced. However, this encapsulated structure makes DNA extraction from the filter cartridge more problematic than that from traditional membrane filters.

There are two major approaches for extracting DNAs from a filter cartridge at present. One of the common approaches is adding lysis buffer directly into a filter cartridge (e.g., Somerville et al. 1989, Sharma et al. 2013, Bryant et al. 2016). In such a protocol, after the addition of lysis buffer into the filter unit, cell lysis is performed inside the filter cartridge. This type of protocol has recently been applied to macrobial environmental DNA studies as well, and showed higher performance than the use of traditional filters (Miya et al. 2016, Spens et al. 2016). In the other approach, the cartridge is broken apart, and then DNA extraction is performed outside the cartridge using a commercial kit (e.g., DNeasy® PowerSoil DNA Isolation kit, Qiagen) which usually includes a bead-beating step (e.g., see video in https://bit.ly/2IKprAS) (Urakawa et al. 2010, Perrine et al. 2017). Although this approach seems to improve the efficiency of DNA extraction, it obviously increases the risk of contamination as well as handling time.

Although the bead-beating step is generally recommended in microbial studies (e.g., Albertsen et al. 2015), the risk of contamination as well as handling time should be reduced if possible. In the present study, I included a manual, intra-cartridge, bead-beating step in a DNA extraction protocol using a DNeasy® Blood & Tissue kit without breaking apart the cartridge (hereafter, “Beads” method), and compared the performance with those of two other protocols that are commonly used in environmental studies: the first one is performing cell lysis inside the filter cartridge using a DNeasy® Blood & Tissue kit without a bead-beating step (hereafter, “NoBeads” method), and the second one is breaking apart the filter cartridge and then performing DNA extraction using a DNeasy® PowerSoil DNA Isolation kit (hereafter, “PowerSoil” method). Specifically, water samples were collected from four different aquatic ecosystems (lake, river, small artificial pond and coastal ecosystems), and DNAs were extracted using the three protocols. Then, the V4 region of the prokaryotic 16S rRNA gene was amplified using 515F-806R primers (Bates et al. 2011, Caporaso et al. 2011). Internal standard DNAs were included in the library preparation process to estimate the number of 16S rRNA genes (internal standard DNA approach; Ushio et al. 2018). The DNA library was sequenced using Illumina MiSeq, and the sequences obtained were analyzed using the amplicon sequence variant (ASV) approach (DADA2; Callahan et al. 2016) and taxa assignments were done using Claidnet (Tanabe and Toju 2013). I found that, 1) the total prokaryotic DNA yields were highest when using the Beads method, 2) The number of ASVs (a proxy for species richness) was highest when using the Beads method, 3) overall community compositions evaluated using the three methods were significantly different among the three methods, and 4) the number of method-specific ASVs was highest when using the Beads method. These results were generally robust across samples from all types of aquatic ecosystems examined.

## Methods

### Preparation of the filter cartridge and beads

For the collection of water samples and on-site filtration, *φ* 0.22-*µ*m Sterivex™ filter cartridges (SVGV010RS, Merck Millipore, Darmstadt, Germany) were used (Fig. 1a-c). When performing field sampling, the filter cartridges were capped with inlet and outlet caps (inlet cap = Luer Fitting VRMP6; outlet cap = Luer Fitting VRSP6, AsOne, Osaka, Japan). The use of caps improved sample handling efficiency and reduced the risk of contamination.

**Figure 1:**
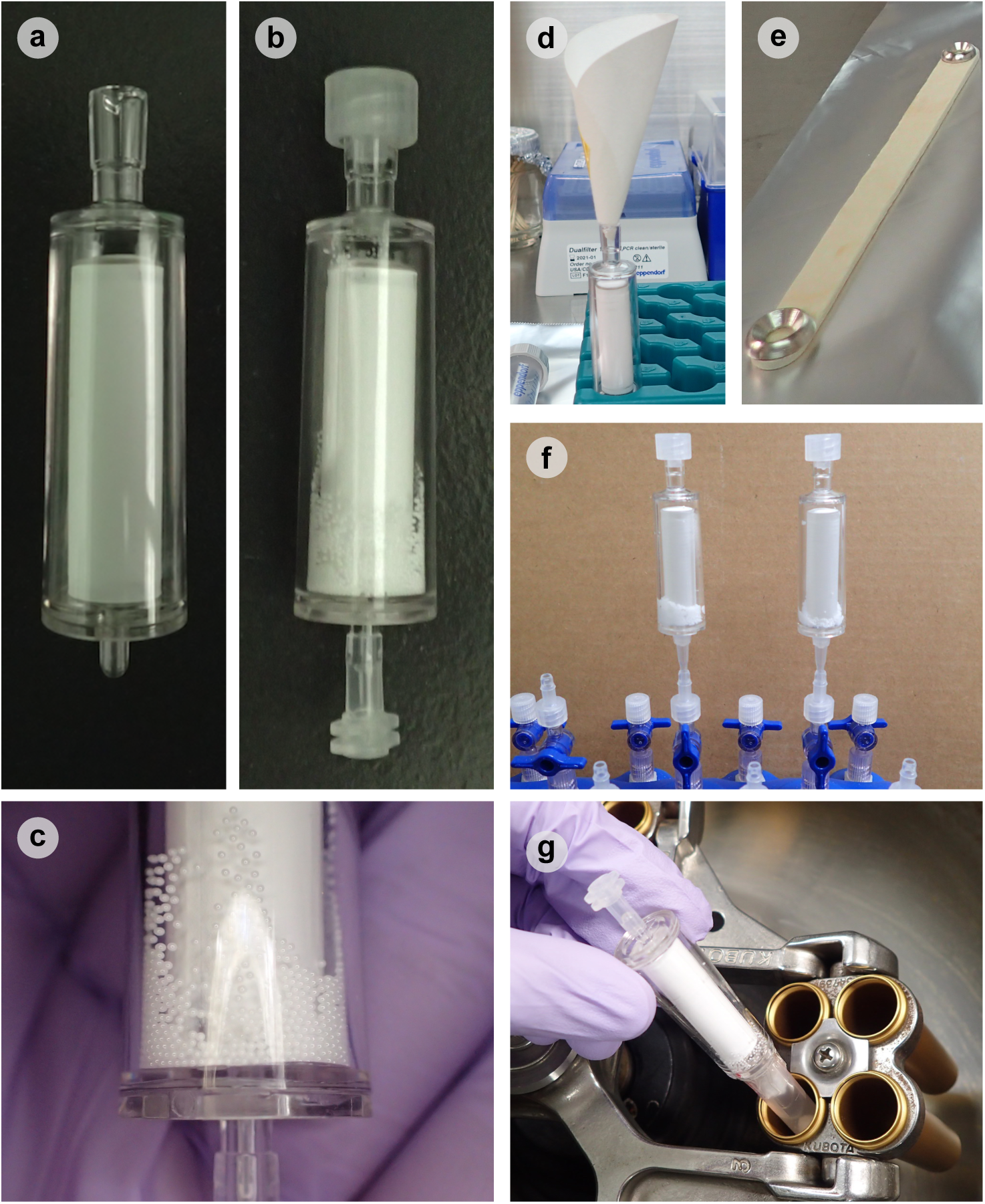
Images of Sterivex™ filter cartridge and zirconia beads. (**a**) Sterivex cartridge filter without zirconia beads. (**b**) Sterivex filter cartridge with inlet/outlet caps (inlet, Luer Fitting VRMP6; outlet, Luer Fitting VRSP6). (**c**) Enlarged image of φ 0.5 mm zirconia beads. (**d**) Sterivex™ filter cartridge with a rolled filter paper (ADVANTEC, 5A 90 mm). (**e**) A hand-made scoop. The scoop is used to add/measure 1 g of zirconia beads. (**f**) RNAlater solution was removed using QIAvac system (Qiagen, Hilden, Germany) before DNA extraction. (**g**) The lysate was transferred into a new 2-ml tube from the inlet of the filter cartridge by centrifugation. A capless 2-ml tube was lightly attached by a clean tape to the outlet of the cartridge before the centrifugation.

For bead-beating, zirconia beads (*φ* 0.5 mm, YTZ-0.5, AsOne, Osaka, Japan) were used. Zirconia beads were purchased separately from the filter cartridges, and they were sterilized under UV light for *ca*. 30 min before use. Then the beads were added into the filter unit in a clean room from the filter unit’s inlet by using a rolled filter paper and a hand-made scoop to avoid static electricity (Fig. 1d,e). After the beads were added into the filter unit, the inlet and outlet were securely closed by inlet/outlet caps. The beads-added, securely-capped filter units were packed and stored in a clean plastic bag until use (e.g., days to months; influences of the storage durations of the beads-added filter units were not explicitly examined in the present study). All processes of the bead preparations were performed in a clean room. Preliminary experiments suggested that the addition of 1 g of the beads improved the yield of total DNA extracted from the filter cartridges (Fig. S1). Based on these preliminary experiments, 1 g of zirconia beads were added to the filter cartridge in this study.

### Study sites and water sampling

Water samplings were performed two times to collect samples from four study sites. For the first sampling, water samples were collected from two sites on 21 February 2017: Lake Biwa (34° 55*′* 45*″* N, 135° 56*′* 41*″* E) and Tenjin River (35° 4*′* 19*″* N, 135° 55*′* 59*″* E) in Shiga prefecture, Japan (Table 1). For the second sampling, water samples were collected from two other sites on 28 December 2018: a small artificial pond in the Center for Ecological Research, Kyoto University (34° 58*′* 18*″* N, 135° 57*′* 33*″* E) and a coastal area in Kobe (34° 38*′* 27*″* N, 135° 13*′* 39*″* E). These sites are hereafter referred to as “Lake”, “River”, “Pond”, and “Sea”, respectively. These sites were selected because they represent contrasting aquatic ecosystems: the lake site represents a relatively stable and mesotrophic aquatic environment, the river site represents a dynamic, relatively clear and oligotrophic aquatic environment, the pond site represents a stable and artificial environment, and the sea site represents a dynamic marine environment. Water samples were collected to test three DNA extraction methods (Fig. 2): 1) a protocol using a filter cartridge with a bead-beating step inside the cartridge, 2) a protocol using a filter cartridge without a bead-beating step and 3) a protocol using a DNeasy® PowerSoil DNA Isolation kit after breaking open a filter cartridge, a common method in microbial ecology studies (e.g., Urakawa et al. 2010, Perrine et al. 2017). Protocols 1) and 2) used a DNeasy® Blood & Tissue Kit (Qiagen, Hilden, Germany). Hereafter, Protocols 1, 2 and 3 are referred to as “Beads” (i.e., with bead-beating), “NoBeads” (i.e., without bead-beating), and “PowerSoil” (i.e., DNeasy® PowerSoil DNA Isolation kit), respectively.

**Table 1:** Study site description.

| Site name | Lake Biwa | Tenjin River | CER Pond | Kobe Sea |
| --- | --- | --- | --- | --- |
| Site type | Lake | River | Pond | Sea |
| Sampline date | 21 Feb 2017 | 21 Feb 2017 | 28 Dec 2018 | 28 Dec 2018 |
| Time | 10:58 | 14:18 | 14:35 | 11:07 |
| Latitude | 35° 4' 19" N | 34° 55' 45" N | 34° 58' 18" N | 34° 38' 27" N |
| Longitude | 135° 55' 59" E | 135° 56' 41" E | 135° 57' 33" E | 135° 13' 39" E |
| Water temperature | 6.1°C | 5.5°C | 10.6°C | 13.9°C |
| Water pH | 8.54 | 8.53 | 7.55 | 7.8 |
| EC (mS/cm) | 0.16 | 0.03 | 0.14 | 4650 |
| Filtered water | 20 ml | 30 ml | 100 ml | 100 ml |

**Figure 2:**
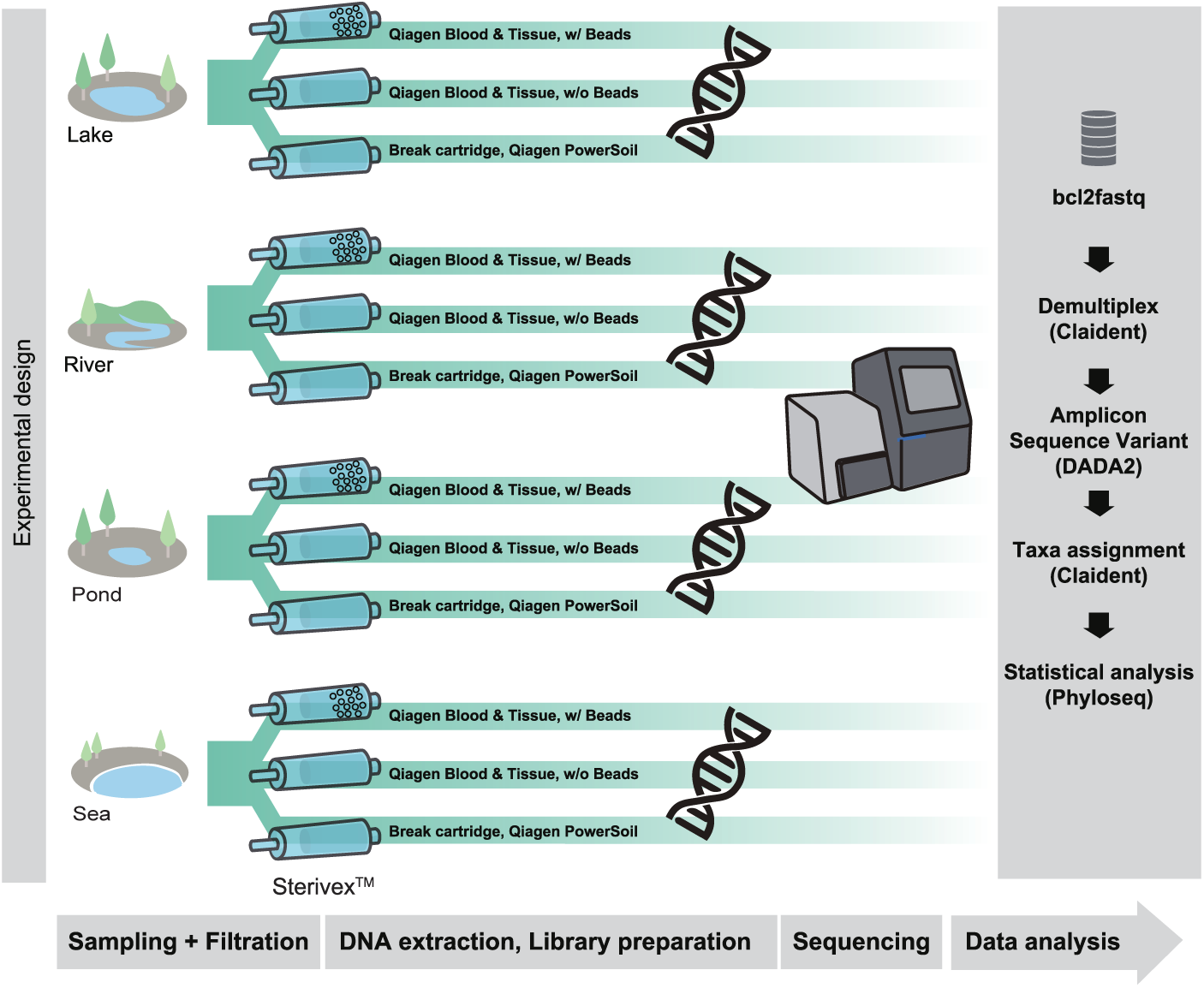
Experimental design of this study. Four research sites (a lake, river, pond and sea) and three DNA extraction methods (Beads, NoBeads, PowerSoil) were examined. Five replications and one field negative control were included for each treatment. After sequencing, bcl2fastq, Claidnet, DADA2 and phyloseq were used to analyze data (see Methods for details).

For the lake, river and pond sites, six 500-ml water samples were collected from six locations of each sampling site (within a *ca*. 100 m diameter area for the lake and river sites, and within a *ca*. 10 m diameter area for the small pond site), and then the six replicates were combined as a 3-L composite water sample in a large plastic bottle. The composite sample was mixed thoroughly to reduce subsample-level variations. For the sea site, an approximately 5-L water sample was collected using a pail that had been sterilized using bleach, and the collected water sample was transferred to a large plastic bottle to mix the water thoroughly and perform water filtration. For each DNA extraction protocol, five subsamples each from the lake, river, pond and sea were obtained by filtering 20, 30, 100 and 100 ml of the pooled water sample, respectively, using a sterile 50-mL syringe (TERUMO, Tokyo, Japan) and a *φ* 0.22-*µm* Sterivex™ filter cartridge with/without zirconia beads inside. The filtered water volumes were adjusted depending on the water sample conditions (e.g., the amount of fine particles that would clog the filter pores). One negative control for each treatment (i.e., a field negative control) was prepared by filtering MilliQ water in the field. In total, 72 water samples were collected as filter cartridges from the four study sites (i.e., four study sites × three DNA extraction protocols×five replicates + twelve field negative controls). After filtration, 2 ml of RNAlater® solution (ThermoFisher Scientific, Waltham, Massachusetts, USA) was added into each filter cartridge. Then, the samples were taken back to the laboratory as soon as possible. During the transport (for up to 2 hours), the samples were kept at 4°C.

### DNA extraction

DNA extraction was performed using either a DNeasy® Blood & Tissue kit or DNeasy® PowerSoil DNA Isolation kit. First, the 2 ml of RNAlater® solution in each filter cartridge was removed from the outlet under vacuum using the QIAvac system (Qiagen, Hilden, Germany), followed by a further wash using 1 ml of MilliQ water. The MilliQ water was also removed from the outlet using the QIAvac system.

For the Beads and NoBeads methods, DNA was extracted from cartridge filters using a DNeasy® Blood & Tissue kit following a protocol described by Miya et al. (2016). Briefly, proteinase-K solution (20 *µ*l), PBS (220 *µ*l) and buffer AL (200 *µ*l) were mixed, and 440 *µ*l of the mixture was added to each filter cartridge. The materials on the cartridge filters were subjected to cell-lysis conditions by incubating the filters on a rotary shaker (15 rpm; DNA oven HI380R, KURABO, Osaka, Japan) at 56°C for 10 min. For the Beads method only, filter cartridges were vigorously shaken (with zirconia beads inside the filter cartridges) for 180 sec (3200 rpm; VM-96A, AS ONE, Osaka, Japan). The incubated and lysed mixture was transferred into a new 2-ml tube from the inlet (not the outlet) of the filter cartridge by centrifugation (3,500 g for 1 min; Fig. 1f). For the Beads method, zirconia beads were removed by collecting the supernatant of the incubated mixture after the centrifugation. The collected DNA was purified using a DNeasy® Blood & Tissue kit following the manufacturer’s protocol. After the purification, DNA was eluted using 100 *µ*l of the elution buffer provided with the kit. Eluted DNAs were stored at −20°C until further processing.

For the PowerSoil method, each cartridge was broken open using sterilized pliers, and the filter was taken from the cartridge and cut into small sections. Then, DNA was extracted from the sections using a DNeasy® PowerSoil DNA Isolation kit. Briefly, the small filter sections were put into a PowerBeads tube, and 60 *µ*l of Solution C1 was added. Then, the tube was incubated at 56°C for 10 min followed by 180 sec of bead beating (3200 rpm). For other processes, I followed the manufacturer’s instructions and DNAs were eluted using 100 *µ*l of Solution C6. Eluted DNAs were stored at −20°C until further processing.

### Paired-end library preparation and sequencing

Prior to the library preparation, work-spaces and equipment were sterilized. Filtered pipet tips were used, and separation of pre- and post-PCR samples was carried out to safeguard against cross-contamination. Four PCR-level negative controls (i.e., with and without internal standard DNAs) were employed for each MiSeq run to monitor contamination during the experiments (Table S1).

The first-round PCR (first PCR) was carried out with a 12-*µ*l reaction volume containing 6.0 *µ*l of 2×KAPA HiFi HotStart ReadyMix (KAPA Biosystems, Wilmington, WA, USA), 0.7 *µ*l of 515F primer, 0.7 *µ*l of 806R primer (updated version described in Earth Microbiome Project [http://www.earthmicrobiome.org/protocols-and-standards/16s/]; each primer (at 5 *µ*M) used in the reaction; with adaptor and six random bases) (Bates et al. 2011, Caporaso et al. 2011), 2.6 *µ*l of sterilized distilled H_2_O, 1.0 *µ*l of DNA template and 1.0 *µ*l of prokaryotic internal standard DNA (for internal standard DNA, see the following subsection Internal standard DNA and quantitative MiSeq sequencing). The thermal cycle profile after an initial 3 min denaturation at 95°C was as follows (35 cycles): denaturation at 98°C for 20 s; annealing at 60°C for 15 s; and extension at 72°C for 30 s, with a final extension at the same temperature for 5 min. We performed triplicate first PCR, and these replicate products were pooled in order to mitigate the PCR dropouts. The pooled first PCR products were purified using AMPure XP (PCR product: AMPure XP beads = 1:0.8; Beckman Coulter, Brea, California, USA). The pooled, purified, and 10-fold diluted first PCR products were used as templates for the second-round PCR.

The second-round PCR (second PCR) was carried out with a 24-*µ*l reaction volume containing 12 *µ*l of 2× KAPA HiFi HotStart ReadyMix, 1.4 *µ*l of each primer (each primer at 5 *µ*M in the reaction volume), 7.2 *µ*l of sterilized distilled H_2_O and 2.0 *µ*l of template. Different combinations of forward and reverse indices were used for different templates (samples) for massively parallel sequencing with MiSeq (Table S1). The thermal cycle profile after an initial 3 min denaturation at 95°C was as follows (12 cycles): denaturation at 98°C for 20 s; annealing at 68°C for 15 s; and extension at 72°C for 15 s, with a final extension at 72°C for 5 min.

Twenty microliters of the indexed second PCR products were mixed, and the combined DNA library was again purified using AMPure XP (PCR product: AMPure XP beads = 1:0.8). Target-sized DNA of the purified library (*ca*. 440 bp) was excised using E-Gel SizeSelect (ThermoFisher Scientific, Waltham, MA, USA). The double-stranded DNA concentration of the library was quantified using a Qubit dsDNA HS assay kit and a Qubit fluorometer (ThermoFisher Scientific, Waltham, MA, USA). The double-stranded DNA concentration of the library was then adjusted using MilliQ water and the DNA was then applied to the MiSeq (Illumina, San Diego, CA, USA). The library which contained the lake and river samples was sequenced in March 2017 using a MiSeq Reagent Kit v2 for 2×250 bp PE, and the library which contained the pond and sea samples was sequenced in February 2019 using a MiSeq Reagent Kit v3 for 2×300 bp PE. Note that samples for other research projects were simultaneously sequenced in the MiSeq runs, and thus not all sequence reads generated were analyzed in the present study (see Table S1 for the number of sequence reads for each run).

### Internal standard DNA and quantitative MiSeq sequencing

Five artificially designed and synthesized internal standard DNAs, which are similar but not identical to the V4 region of any existing prokaryotic 16s rRNA, were included in the library preparation process to estimate the number of prokaryotic DNA copies (i.e., quantitative MiSeq sequencing; see Ushio et al. 2018). They were designed to have the same primer-binding regions as those of known existing prokaryotes and conserved regions in the insert region (Table S2). Variable regions in the insert region were replaced with random bases so that no known existing prokaryotic sequences had the same sequences as the standard sequences. The numbers of standard DNA copies were adjusted appropriately to obtain a linear regression line between standard DNA copy numbers and their sequence reads from each sample.

Ushio et al. (2018) showed that the use of standard DNAs in MiSeq sequencing provides reasonable estimates of the DNA quantity in environmental samples when analyzing fish environmental DNA. However, it should be noted that it corrects neither for sequence-specific amplification efficiency, nor for species-specific DNA extraction efficiency. In other words, the method assumes similar amplification efficiencies across sequences, and similar DNA extraction efficiencies across microbial species, which is apparently not valid for complex environmental samples, and thus we need careful interpretations of results obtained using that method.

### Sequence data processing

All scripts to process raw sequences are available at https://github.com/ong8181/micDNA-beads. The raw MiSeq data were converted into FASTQ files using the bcl2fastq program provided by Illumina (bcl2fastq v2.18). The FASTQ files were then demultiplexed using the command implemented in Claident (http://www.claident.org; Tanabe and Toju 2013). I adopted this process rather than using FASTQ files demultiplexed by the Illumina MiSeq default program in order to remove sequences whose 8-mer index positions included nucleotides with low quality scores (i.e., Q-score *<* 30).

Demultiplexed FASTQ files were analyzed using the Amplicon Sequence Variant (ASV) method implemented in DADA2 v1.10.1 (Callahan et al. 2016). I chose to use the ASV approach implemented in DADA2 rather than using a more common and traditional OTU approach implemented in e.g., QIIME (Caporaso et al. 2010). This is because ASV methods, including DADA2, are able to infer the biological sequences in the sample prior to the introduction of amplification and sequencing errors, and are superior when one may wish to distinguish between sequence variants differing by as little as one nucleotide (Callahan et al. 2016). Considering their fine resolution, reusability, reproducibility and comprehensiveness, ASV approaches have recently been recommended as a primary choice for microbial marker-gene analysis (e.g., Callahan et al. 2017, Note that ASV methods are also a primary choise in QIIME2, https://qiime2.org/). At the quality filtering process, forward and reverse sequences were trimmed at the length of 215 and 160 based on the visual inspection of Q-score distribution, respectively, using DADA2::filterAndTrim() function. The two MiSeq runs were performed using a different sequencing kit (i.e., v2 kit for 2×250 bp PE and v3 kit for 2×300 bp PE), but the sequence lengths were trimmed at the same length at this step. Error rates were learned using DADA2::learnErrors() function (MAX_CONSIST option was set as 20). Then, sequences were dereplicated, error-corrected, and merged to produce an ASV-sample matrix. Chimeric sequences were removed using the DADA2::removeBimeraDenove() function.

Taxonomic identification was performed for ASVs inferred using DADA2 based on the query-centric auto-*k*- nearest-neighbor (QCauto) method (Tanabe and Toju 2013) and subsequent taxonomic assignment with the lowest common ancestor algorithm (Huson et al. 2007) using “semiall” database (i.e., database includes all sequences prokaryotes and eukaryotes except vertebrates, *Caenorhabditis* and *Drosophila*) and clidentseq and classigntax commands implemented in Claident v0.2.2018.05.29. Because the QCauto method requires at least two sequences from a single microbial taxon, only internal standard DNAs were separately identified using BLAST (Camacho et al. 2009).

### Estimations of DNA copy numbers

For all analyses in this subsection, the free statistical environment R 3.5.2 was used (R Core Team 2018). Rarefying sequence reads is a common approach to evaluate microbial diversity. I examined rarefaction curves and found that the sequencing captured most of the prokaryotic diversity (Fig. S2). Also, a previous study suggested that simply rarefying microbial sequence data is inadmissible (McMurdie and Holmes 2014). Another important point is that my analyses involve conversion of sequence reads to DNA copy numbers and thus are different from other commonly used analyses in microbiome studies. Considering these conditions and the previous study, sequence reads were subjected to most of the downstream analysis without performing further corrections using rarefaction or other approaches in the present study.

The procedure to estimate DNA copy numbers consisted of two parts following the previous study (Ushio et al. 2018): (1) linear regression analysis to examine the relationship between sequence reads and the copy numbers of the internal standard DNAs for each sample, and (2) the conversion of sequence reads of non-standard prokaryotic DNAs to estimate the copy numbers using the result of the linear regression for each sample.

Linear regressions were used to examine how many sequence reads were generated from one microbial DNA copy through the library preparation process. Note that a linear regression analysis between sequence reads and standard DNAs was performed for each sample and the intercept was set as zero. The regression equation was: MiSeq sequence reads = regression slope×the number of standard DNA copies [*/µ*l]. The number of linear regressions performed was 76 (= the number of microbial DNA samples + PCR negative control with standard DNAs), and thus 76 regression slopes were estimated in total.

The sequence reads of non-standard prokaryotic DNAs were converted to calculated copy numbers using sample-specific regression slopes estimated by the above regression analysis. The number of non-standard prokaryotic DNA copies was estimated by dividing the number of MiSeq sequence reads by a sample-specific regression slope (i.e., the number of DNA copies = MiSeq sequence reads*/*regression slope). A previous study demonstrated that these procedures provide a reasonable estimation of DNA copy numbers using high-throughput sequencing (Ushio et al. 2018).

### Statistical analyses

All scripts in this section are available at https://github.com/ong8181/micDNA-beads. Visualization and basic statistical analyses were performed using the phyloseq v1.26.1 package of R (McMurdie and Holmes 2013). First, sample metadata, ASV-sample matrix (equivalent to classical “OTU table”), and taxa information were imported to a phyloseq object using phyloseq::phyloseq() function. Visualization and basic statistical analyses consisted of four parts: 1) total prokaryotic DNA copy numbers and the number of ASVs were compared using Tukey-HSD, 2) broad-scale phylogenetic compositions were visualized using a barplot, 3) community compositions were compared using non-metric dimensional scaling (NMDS) and 4) method-specific microbial ASVs (i.e., ASVs frequently detected with a certain method, but rarely detected with the other methods) were identified. During the processes of the statistical analyses and visualizations, functionalities implemented in R packages of “vegan” (Oksanen et al. 2008), “tidyverse” (Wickham 2017), “ggplot2” (Wickham 2009), “cowplot” (Wilke 2017) and “ggsci” (Xiao 2018) were also used. Detailed information on the package version is available in “00_SessionInfo_original” folder at https://github.com/ong8181/micDNA-beads.

### Sequence data and code availability

DDBJ Accession numbers of the BioProject of the DNA sequences analyzed in the present study are DRA006959. Scripts to implement sequence processing, statistical analyses and visualization were available at https://github.com/ong8181/micDNA-beads.

## Results

### Sequence reads and the copy numbers of standard DNAs

In total, MiSeq run and DADA2 processing generated 803,841 high-quality, merged reads, excluding internal standard DNAs (*mean* = 12, 945 reads, *S.D.* = 7, 465; Table S1). In the first MiSeq run, 11–49% and 0.2–1.9% of sequence reads were from non-standard microbial DNAs for field samples and field negative controls, respectively. In the second MiSeq run, 68–96% and 1.7–10% of sequence reads were from non-standard microbial DNAs for field samples and field negative controls, respectively.

The relationships between sequence reads and the copy numbers of standard DNAs were examined by linear regression analysis (Fig. 3). Within each sample, the sequence reads linearly and positively correlated with the copy numbers of standard DNAs (Fig. 3a). Variations explained by the linear regressions were over 0.9 for most samples (Fig. 3b). These results suggested that the number of sequence reads was generally proportional to the number of copies of DNA initially added to the first PCR reaction within a single sample. The slopes of the linear regressions varied depending on the sample, suggesting that the amplification efficiency depended on the individual sample. These sample-specific slopes were used to convert sequence reads to the copy numbers of microbial ASVs.

**Figure 3:**
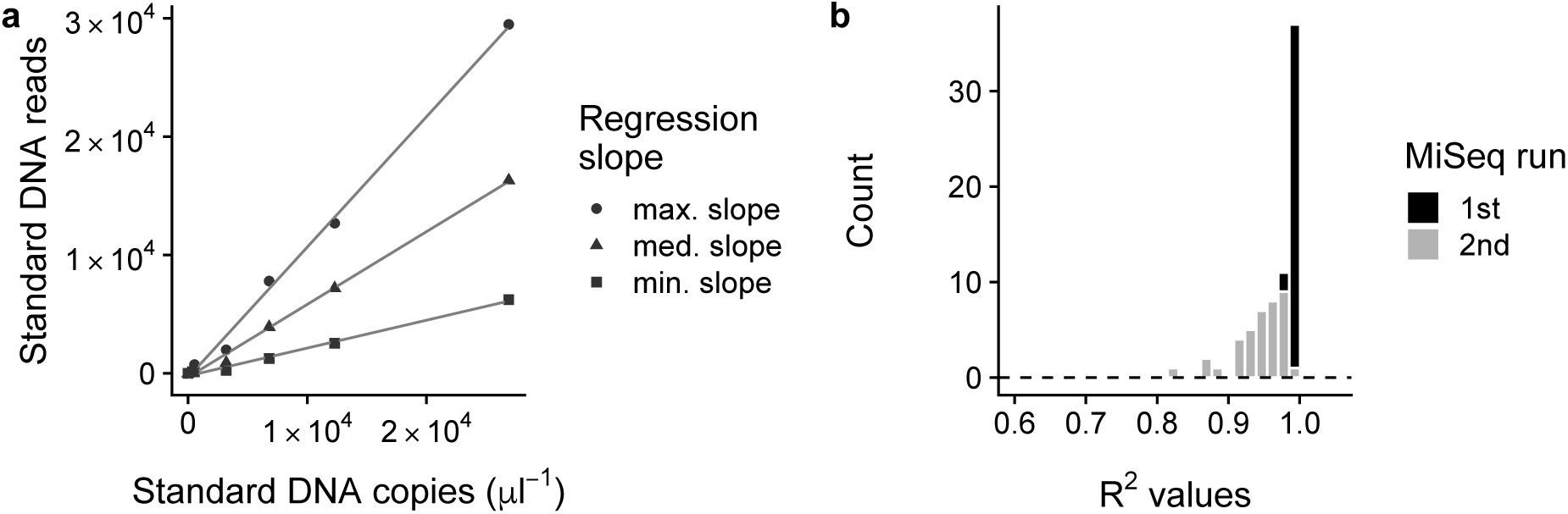
The relationships between sequence reads and the copy numbers of standard DNAs. (**a**) Examples of the relationships between sequence reads and the copy numbers of standard DNAs. The linear regressions with the maximum, median and minimum slopes of the first MiSeq run are shown. (**b**) Distribution of R^2^ values of the linear regression for the first and second MiSeq runs.

Levels of contaminations were examined using the field negative controls. DNA copy numbers detected in the field negative controls were less than 0.07% of mean DNA copy numbers of field positive samples (Fig. S3), suggesting that there was only a small amount of contamination during the library preparation process. In addition, there were no qualitative differences in DNA copy numbers or phyla detected among the DNA extraction methods (Fig. S3).

### Total prokaryotic DNA quantity and the number of ASVs

The estimated copy numbers of prokaryotic ASVs were summed to estimate the total numbers of prokaryotic DNAs (Fig. 4a). The total prokaryotic DNA copy numbers obtained by the Beads method were significantly higher than those obtained by the NoBeads and PowerSoil methods for the lake, pond and sea samples (Fig. 4a). This was consistent with NanoDrop quantification, which showed that the Beads method generally yielded higher amounts of DNA than the other two methods for (Fig. S4). While there was no significant difference in the total amounts of prokaryotic DNA for the river samples, the mean values of the amount of prokaryotic DNAs obtained by the Beads and NoBeads methods were higher than that obtained by the PowerSoil method, which is also consistent with the NanoDrop results (Figs. 4a and S4). There were no significant differences in the numbers of ASVs (an index of species richness) obtained using the three different extraction methods when all ASVs were taken into account (Fig. 4b).

**Figure 4:**
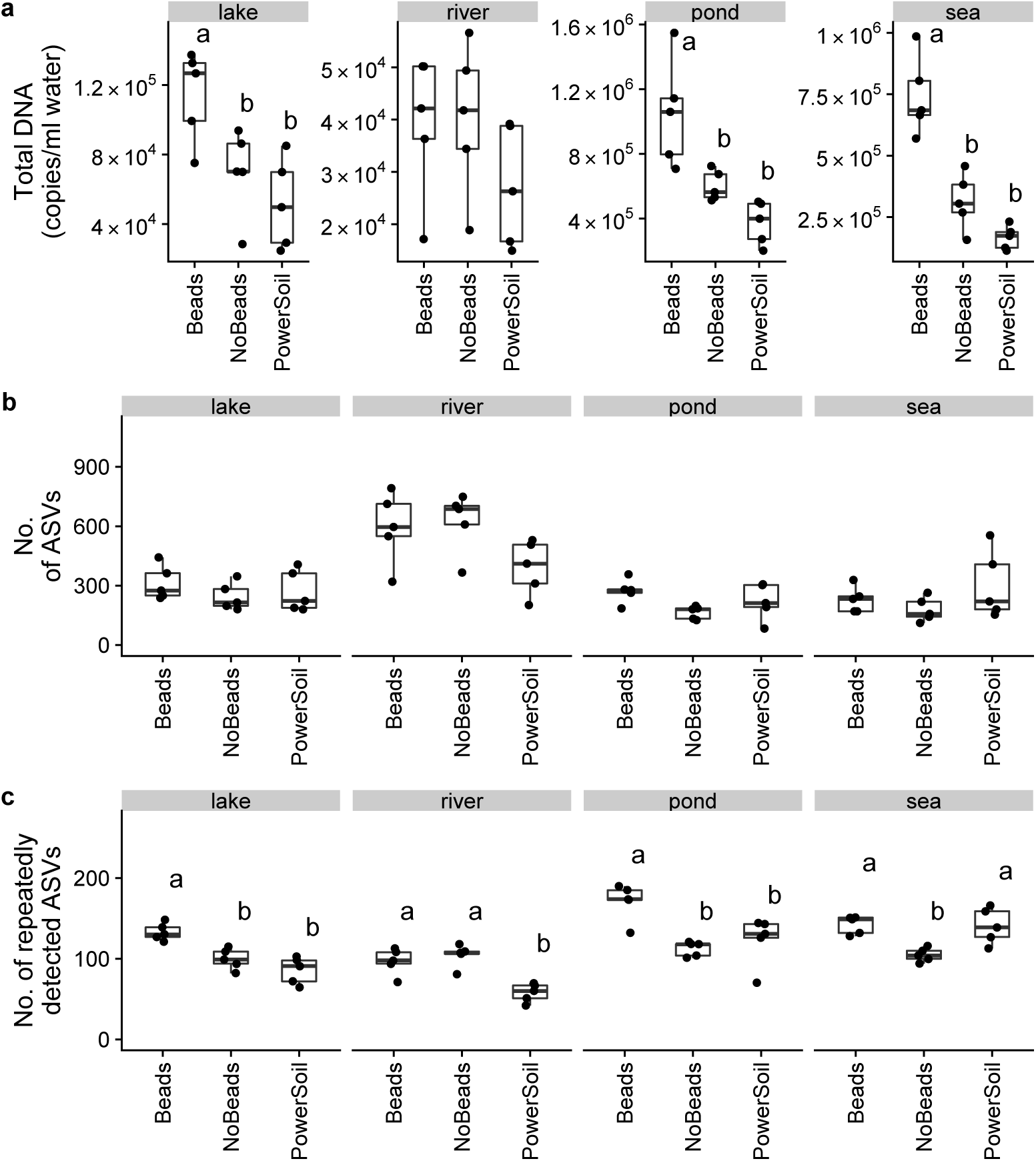
Total prokaryotic DNA copy numbers and the number of ASVs for each treatment. (**a**) Total prokaryotic DNA copy numbers calculated by summing the estimated DNA copy numbers of all prokaryotic ASVs. (**b**) The number of all ASVs. (**c**) The number of repeatedly detected ASVs. Different letters indicate significant differences between treatments (P < 0.05).

Rare ASVs (e.g., detected only once among the five replicates in each method) were dominant among the detected ASVs (Table 2). On average, 650, 1948, 284 and 409 ASVs were detected only once among the five replicates for lake, river, pond and sea samples, respectively (Table 2). The numbers of ASVs that were detected in all of the five replicates were on average 52, 32, 80 and 82 in the lake, river, pond and sea samples, respectively (Table 2). When only frequently detected ASVs (i.e., ASVs detected at least three times among the five replicates) were considered, the number of ASVs detected in the Beads method was significantly higher than that in the other two methods for the lake and pond samples (Fig. 4c; for lake samples, 133, 100 and 86 ASVs for the Beads, NoBeads and PowerSoil methods, respectively; for pond samples, 170, 112 and 123 ASVs for the Beads, NoBeads and PowerSoil methods, respectively). The numbers of ASVs detected in the Beads and NoBeads methods were significantly higher than that in the PowerSoil method for the river samples (Fig. 4c; 96, 104 and 58 ASVs for the Beads, NoBeads and PowerSoil methods), while the numbers of ASVs detected in the Beads and PowerSoil methods were significantly higher than that in the NoBeads method for the sea samples (Fig. 4c; 142, 104 and 141 ASVs for the Beads, NoBeads and PowerSoil methods).

**Table 2:** The number of ASVs detected in the five replicates in each method and site. Percentage indicates the proportion of estimated DNA copy numbers of the detected frequency category

| Site | Method | Detection frequency | 1 | 2 | 3 | 4 | 5 |
| --- | --- | --- | --- | --- | --- | --- | --- |
| Lake | Beads |  | 684 | 110 | 51 | 43 | 68 |
|  |  |  | 9.8% | 5.7% | 4.6% | 8.4% | 71.5% |
| Lake | NoBeads |  | 555 | 85 | 40 | 36 | 47 |
|  |  |  | 11.0% | 6.1% | 6.9% | 9.4% | 66.6% |
| Lake | PowerSoil |  | 711 | 110 | 44 | 23 | 41 |
|  |  |  | 16.3% | 8.3% | 8.8% | 7.8% | 58.9% |
| River | Beads |  | 2,157 | 165 | 50 | 36 | 38 |
|  |  |  | 47.0% | 10.4% | 7.0% | 9.6% | 26.0% |
| River | NoBeads |  | 2,211 | 191 | 64 | 36 | 37 |
|  |  |  | 44.0% | 12.2% | 9.6% | 9.1% | 25.0% |
| River | Powersoil |  | 1,475 | 98 | 36 | 18 | 22 |
|  |  |  | 48.1% | 12.2% | 8.4% | 7.6% | 23.7% |
| Pond | Beads |  | 341 | 84 | 41 | 39 | 115 |
|  |  |  | 2.5% | 1.4% | 1.4% | 2.5% | 92.2% |
| Pond | No Beads |  | 175 | 43 | 36 | 21 | 74 |
|  |  |  | 1.7% | 1.0% | 2.6% | 1.5% | 93.1% |
| Pond | PowerSoil |  | 335 | 74 | 53 | 50 | 51 |
|  |  |  | 2.2% | 1.9% | 3.1% | 8.0% | 84.8% |
| Sea | Beads |  | 327 | 55 | 33 | 38 | 92 |
|  |  |  | 2.1% | 1.4% | 1.8% | 3.7% | 91.0% |
| Sea | No Beads |  | 269 | 51 | 27 | 22 | 71 |
|  |  |  | 2.9% | 2.2% | 2.5% | 3.4% | 89.0% |
| Sea | PowerSoil |  | 632 | 90 | 54 | 33 | 82 |
|  |  |  | 3.6% | 2.4% | 3.7% | 3.5% | 86.8% |

Although a common data preprocessing, rarefaction, was not performed in the present study (see “*Estimations of DNA copy numbers*” subsection in Methods), the effect of the sequence filtering process was briefly investigated to facilitate the comparison with previous studies. In the analysis, the number of ASVs after removing ASVs with a small DNA copy number was calculated (Fig. S5). This process is equivalent to rarefaction (or, the removal of rare OTUs*/*ASVs) because it removed ASVs with a small copy number, but did not affect the estimated DNA copy numbers of dominant ASVs. The result showed that the pattern of the number of ASVs was not changed compared with the result without the filtering process.

### Prokaryotic community composition

Broad-scale community compositions are shown in Fig. 5. Actinobacteria, Bacteroidetes, Proteobacteria and Verrucomicrobia were dominant phyla in the lake samples, and Bacteroidetes and Proteobacteria were dominant phyla in the river samples (Fig. 5). Proteobacteria was a dominant phylum in the pond samples, and Bacteroidetes, Euryarchaeota and Proteobacteria were dominant phyla in the sea samples (Fig. 5). While the dominant phyla detected were similar among the methods, there were significant differences in overall community composition as determined by nonmetric dimensional scaling (NMDS) and PERMANOVA (only frequently detected ASVs [= ASVs detected at least three times among the five replicates] were considered; *P <* 0.01 for all sites; Fig. 6; for all ASV results, see Fig. S6). On the other hand, inter-site differences in the community compositions were clearly detected by the all extraction methods (Fig. S6a,b).

**Figure 5:**
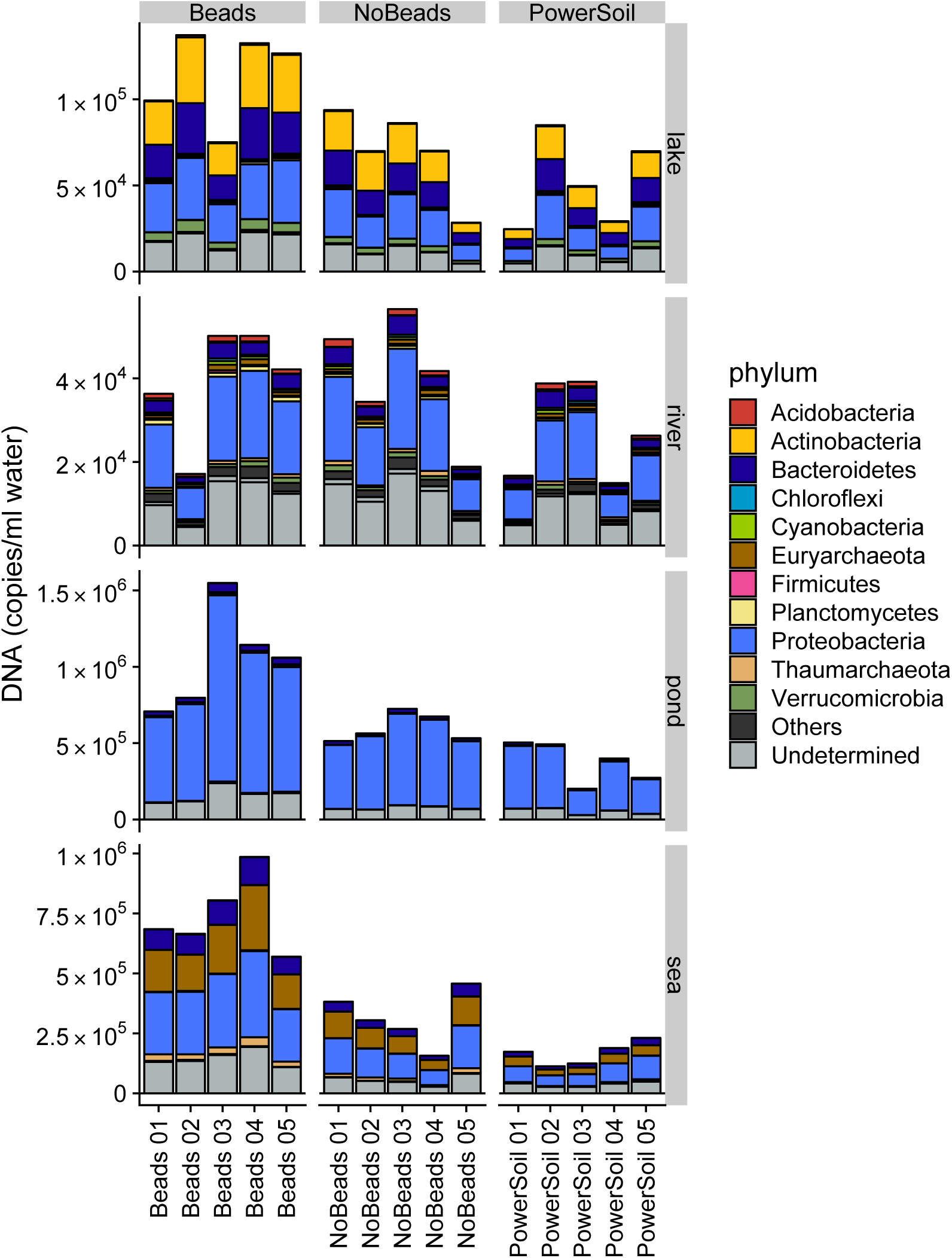
Barplot of detected prokaryotic phyla. Different colours indicate different prokaryotic phyla.

**Figure 6:**
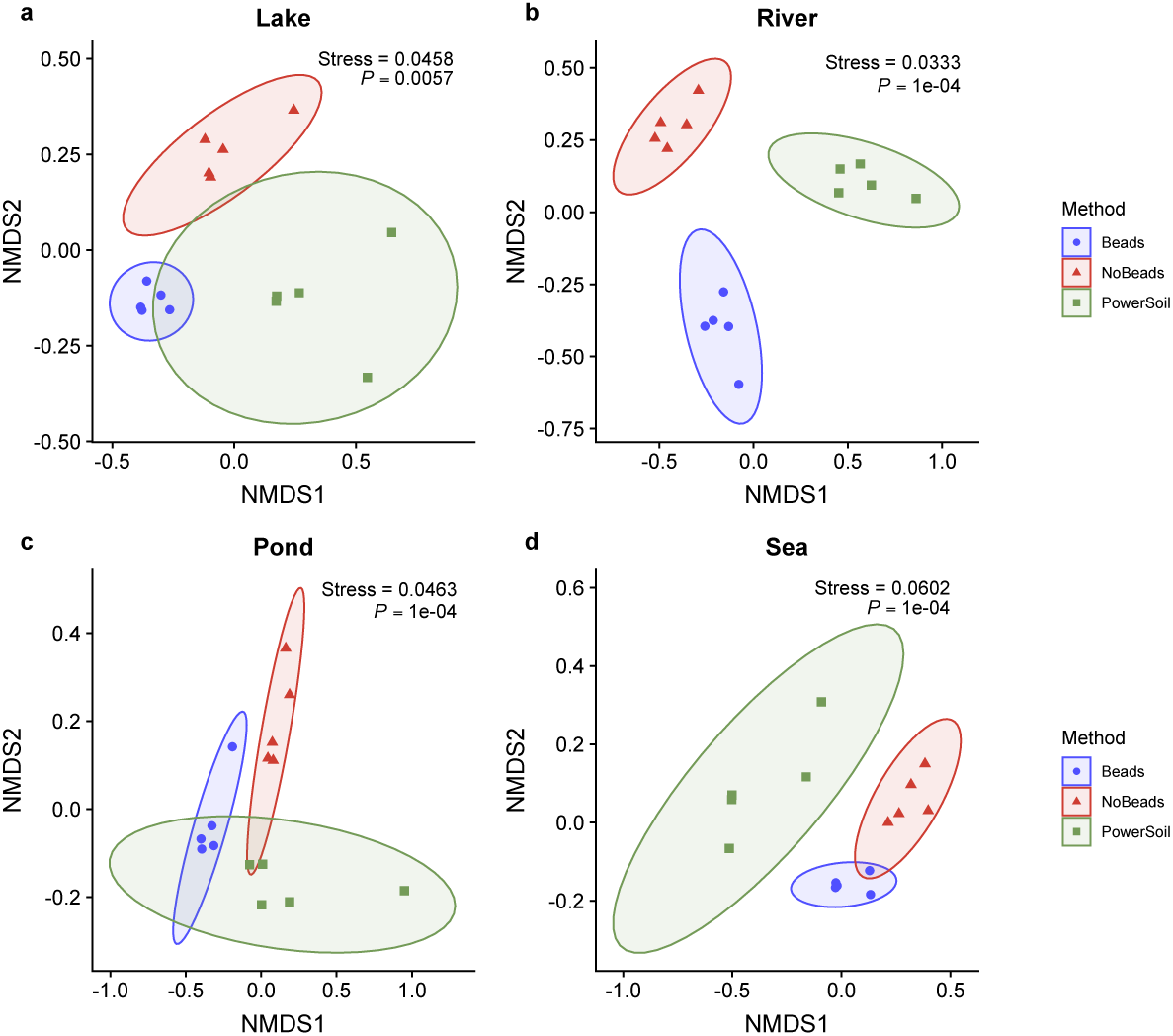
Nonmetric dimensional scaling (NMDS) of the prokaryotic community composition. Only repeatedly detected ASVs were used. NMDS for (**a**) lake, (**b**) river, (**c**) pond and (**d**) sea samples. “P” indicates the significance of the difference among methods on the overall community composition. “Stress” indicates the stress value for each NMDS.

### Method-specific ASVs

In addition to the basic statistical analyses described above, prokaryotic ASVs that were specific to a certain method were investigated (Fig. 7 and Table S3). For this analysis, method-specific ASVs were defined as follows: ASVs that were detected by a specific method at least four times among five replicates, but were detected by the other methods at most two times (FASTA files for the method-specific ASVs are available at https://github.com/ong8181/micDNA-beads/00_fasta/). In the lake samples, 13, 5, and 3 ASVs were identified as Beads-, NoBeads-, and PowerSoil-specific ASVs, respectively. Thus, the Beads method detected the highest number of method-specific ASVs. Among the Beads-specific ASVs in the lake samples, Bacteroidetes (e.g., *Ohtaekwangia*) and Proteobacteria (e.g., Sphingomonadaceae) were detected (Fig. 7 and Table S3). In the river samples, 13, 13, and 1 ASVs were identified as Beads-, NoBeads-, and PowerSoil-specific ASVs, respectively. The Beads and NoBeads methods detected higher numbers of method-specific ASVs than the PowerSoil method. Among the method-specific ASVs in the river samples, Bacteroidetes (*Flavobacterium*), Nitrospirae (*Nitrospira*) and Proteobacteria as well as Crenarchaeota (*Fervidicoccus*) were detected by the Beads method more efficiently than by the other methods. The NoBeads method detected ASVs belonging to Proteobacteria, Firmicutes and Nitropirae more efficiently than the other two methods. In the pond samples, 22, 1, and 3 ASVs were identified as Beads-, NoBeads-, and PowerSoil-specific ASVs, respectively. The Beads method detected several genera belonging to Bacteroidetes, Fusobacteria, Proteobacteria and Verrucomicrobia as well as Thaumarchaeota more efficiently than the other methods. In the sea samples, 10, 1, 8 ASVs were identified as Beads-, NoBeads-, and PowerSoil-specific ASVs, respectively. The Beads method detected several genera belong to Actinobacteria, Bacteroidetes, Proteobacteria and Euryarchaeota more efficiently than the other two methods.

**Figure 7:**
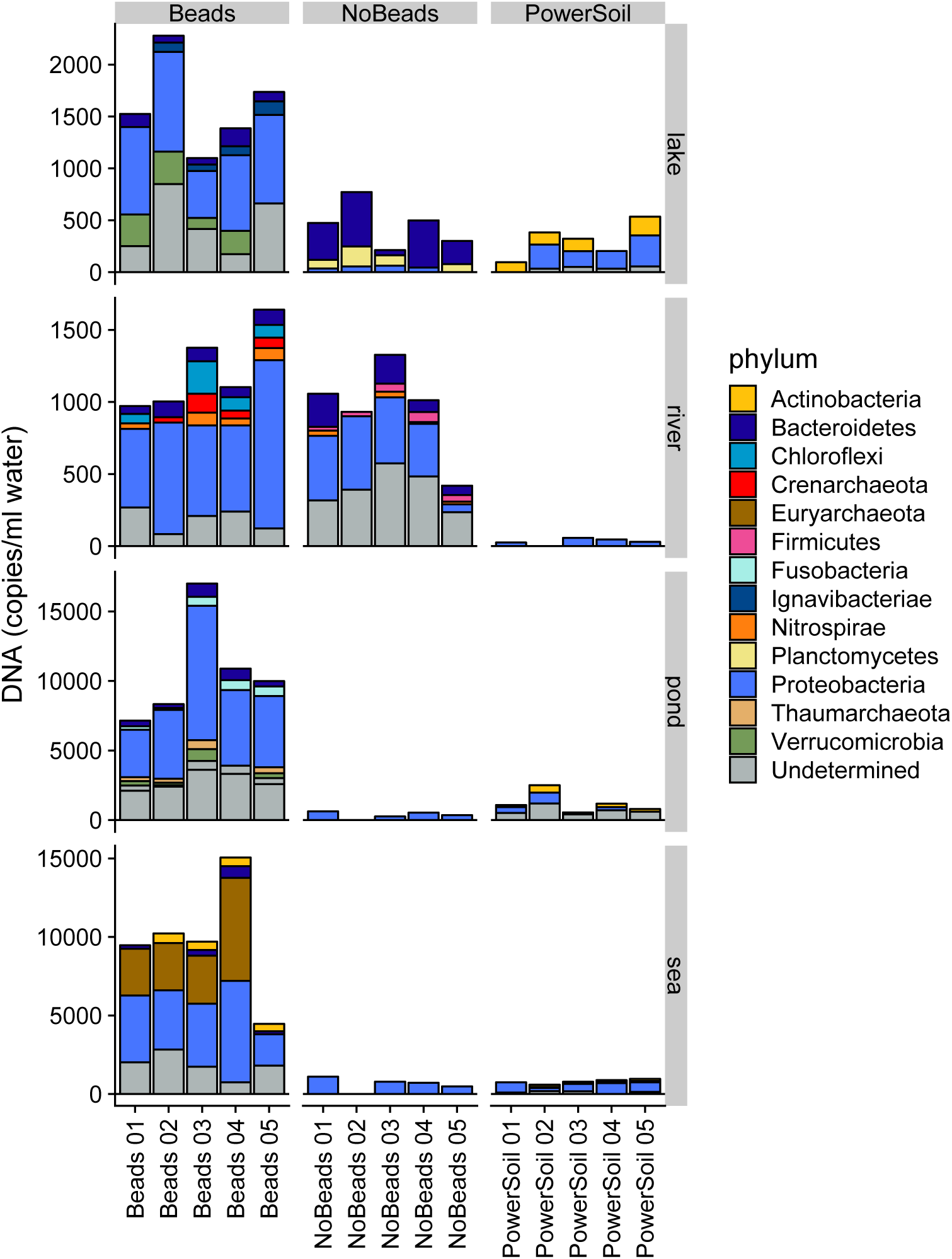
Barplot of method-specific ASVs. Different colours indicate different prokaryotic phyla specific to each DNA extraction method. For the definition of “method-specific ASV”, see the main text.

Although the above-defined criterion was qualitative (i.e., detection frequency rather than estimated copy numbers was used), a more quantitative criterion also showed a similar pattern (Fig. S7). For the lake, pond and sea samples, the Beads method detected a higher number of method-specific ASVs than the other two methods, while for the river samples, the Beads and NoBeads methods detected higher numbers of method-specific ASVs than the PowerSoil method.

## Discussion

### Sequence reads and the copy numbers of standard DNAs

The linear relationships observed between the number of sequence reads and the amount of internal standard DNAs suggested that, within a single sample, the sequence reads were proportional to the copy numbers of DNA. This result suggested that the conversion of sequence reads using a sample-specific regression slope would provide an estimation of DNA copy numbers in the DNA extract (i.e., estimated DNA copy number = sequence reads/sample-specific regression slope). This is consistent with a previous study that showed that this procedure provides a reasonable estimate of multispecies DNA copy numbers for fish environmental DNA (Ushio et al. 2018).

The variations in the regression slopes suggested that there was variation in the loss of PCR products during the library preparation process (e.g., purification step). Another possible source for the variation is that the degree of PCR inhibition is different among samples even though water samples are often considered to contain lower levels of PCR inhibitors than soil samples. Regression slopes could be an index of PCR inhibition if the concentration of DNA (or PCR product) is not normalized. However, this is not the case in my study because the concentration of DNA (or PCR product) was normalized during the purification step using AMPure XP. Therefore, the sources of the variation in the regression slope could not be distinguished in the present study.

### Total prokaryotic DNA quantity and the number of ASVs

Total prokaryotic DNA quantities, estimated by the sum of prokaryotic sequence reads and sample-specific regression slopes, were highest with the Beads method, suggesting that the inclusion of the bead-beating step within the filter cartridge was effective for the extraction of prokaryotic DNAs (Fig. 4), a conclusion which was also supported by the NanoDrop quantification results (Fig. S4). This finding is consistent with other studies that showed that a bead-beating step improved DNA yield when using membrane filters (e.g., Albertsen et al. 2015 and references therein). On the other hand, the PowerSoil method did not improve the DNA yield even though it also included a bead-beating step. This may have been due to the stricter purification step of the PowerSoil® DNA Isolation kit, as it is designed to remove PCR inhibitors (e.g., humic substances) found in soils (see Schrader et al. 2012, Eichmiller et al. 2016). As a result, the DNA yield of the PowerSoil method was relatively low, but the amplification efficiency and/or purity of the extracted DNA could be relatively higher, as shown in other studies (Eichmiller et al. 2016). However, because water samples generally contain smaller amounts of PCR inhibitors than soil samples, the Beads method using a DNeasy® Blood & Tissue kit may be a better choice considering the recovery efficiency of the total DNAs. The recently introduced DNeasy® PowerSoil Pro DNA Isolation kit (Qiagen, Germany), which was not available at the time of the first MiSeq run of the present study, might improve DNA yield as well as DNA purity compared with the DNeasy® PowerSoil DNA Isolation kit. However, like with the DNeasy® PowerSoil DNA Isolation kit, the filter cartridge needs to be broken apart if the PowerSoil® Pro DNA Isolation kit is used, and this will increase the risk of contamination and the handling time.

In terms of the number of total ASVs, there were no significant differences among the methods for either type of samples. However, the numbers of repeatedly detected ASVs were significantly different, and again suggested that beads-beating is an effective method to increase the number of recovered prokaryotic ASVs (Fig. 4c). Also, this result was not affected by filtering ASVs with a small DNA copy number (Fig. S5).

### Prokaryotic community composition and method-specific ASVs

Although there were no clear differences in the coarse-scale community composition (Fig. 5), NMDS of the repeatedly detected ASVs suggested that overall prokaryotic community compositions were significantly different among the DNA extraction methods for all sites (Fig. 6). This suggests that the DNA extraction method would have impacts on the characterization of the overall microbial community composition. Interestingly, the variations in community composition revealed by the Beads or NoBeads methods were smaller than those revealed by the PowerSoil method (Figs. 6 and S6b). Also, the proportion of ASVs detected in all of the five replicates was higher in the Beads or NoBeads method than in the PowerSoil method in all sites (Table 2). These would suggest that the Beads or NoBeads method provided more reproducible results for microbial DNA extraction. The increase in the handling time and steps during the PowerSoil method may contribute to the larger variation found for the PowerSoil method, and the higher DNA yields and higher numbers of ASVs obtained by the Beads method would contribute to the differences in the community composition.

The results of method-specific ASV analyses were generally consistent with the other results in the present study. In general, the number of method-specific ASVs obtained with the Beads method was higher than the number obtained with the NoBeads and PowerSoil method, and the numbers of DNA copies of the method-specific ASVs were also higher with the Beads method than with the other two methods (Fig. 7). This tendency was especially clear for the lake, pond and sea samples.

The results of method-specific ASV analyses were also helpful for inferring a cause of the increased number of ASVs with the Beads method because the method-specific ASVs contributed to the differences in the number of ASVs at least to some extent (i.e., the number of method-specific ASVs/repeatedly detected ASVs of the Beads, NoBeads and PowerSoil methods are summarized as follows: Lake, 13/133, 5/100, 3/86; River, 13/97, 13/104, 1/58; Pond, 22/170, 1/112, 3/123; Sea, 10/141, 1/105, 8/141; see also the Results section). The phyla of the method-specific ASVs of the river, pond and sea samples were likely reflections of the broad-scale community composition (Figs. 5 and 7), suggesting that the cause of the increased numbers of ASVs was not an increased number of ASVs of a certain microbial group, at least at phylum-level. On the other hand, the phyla of the method-specific ASVs of the lake samples were different from those of the broad-scale community composition (Figs. 5 and 7). Proteobacteria (family Sphingomonadaceae) and Ignavibacteriae (genus *Ignavibacterium*) were more frequently detected as a Beads-method-specific phylum (Fig. 7 and Table S3), suggesting that the Beads-method more efficiently extracted DNAs of these microbial groups. Understanding the chemical structure of microbial cell walls would help to reveal more detailed mechanisms of the increased numbers of ASVs obtained with the Beads method.

These results suggested that the inclusion of the intra-cartridge beads-beating step was effective, not only for DNA recovery, but also for the detection of prokaryotic species (Albertsen et al. 2015). It should be noted, however, that, for macrobial environmental DNA, the bead-beating step might decrease the detection efficiency as it is likely that macrobial environmental DNA is highly labile and thus a DNA extraction protocol without bead-beating would be more efficient (Miya et al. 2016, Spens et al. 2016).

## Conclusion

In the present study, I have shown that the inclusion of a bead-beating step inside the filter cartridge improved the yield of DNA recovered from water samples collected from freshwater and marine ecosystems and also the number of prokaryotic ASVs detected compared with two other common approaches used in microbial ecology studies. Performing the bead-beating step in the filter cartridge causes no dramatic increase in either handling time or processing cost. Therefore, this approach has the potential to become one of the major options when one is interested in collecting aquatic microbial samples from the field.

## Acknowledgements

I would like to thank Asako Kawai for assistance in the sampling and experiments, Hiroki Yamanaka for providing an opportunity to use Illumina MiSeq, Shunsuke Matsuoka for assistance in the field sampling, and Ai Matsuda for help in the figure editing. This research was supported by PRESTO (JPMJPR16O2) from the Japan Science and Technology Agency (JST).

