## Supporting information for "Use of a filter cartridge combined with intra-cartridge bead beating improves detection of microbial DNA from water samples"

*Masayuki Ushio*

### Contents:


- **Figure S1**| The relationship between DNA yield quantified by NanoDrop and bead amount
- **Figure S2**| Rarefaction curve of sequence reads of each sample
- **Figure S3**| Prokaryotic phyla detected in the field negative controls
- **Figure S4**| The relationship between DNA yield quantified by NanoDrop and DNA extraction method
- **Figure S5**| The number of ASVs with more than 100 copies/ml water
- **Figure S6**| Non-metric dimensional scaling (NMDS) for ASVs from all study sites
- **Figure S7**| Method-specific ASVs for each DNA extraction method based on an alternative, quantitative criterion
- **Table S1**| Sample metadata and summary for DADA2 sequence processing
- **Table S2**| Prokaryote standard DNA sequences including primer regions
- **Table S3**| Taxa assignments of method-specific ASVs

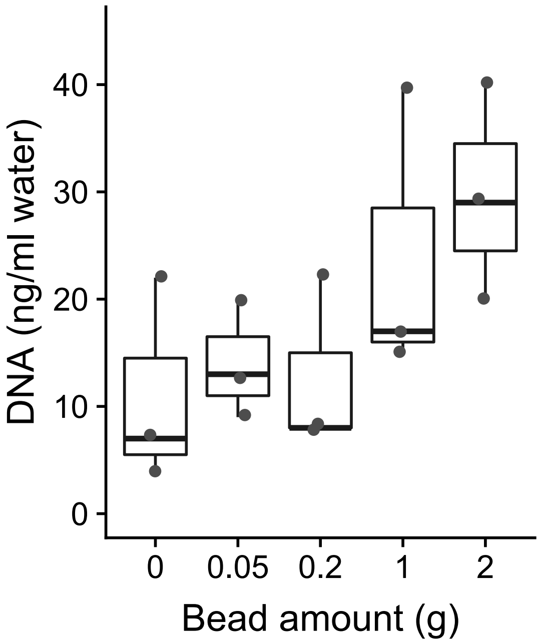


**Figure S1**| The relationship between DNA yield quantified by NanoDrop and bead amount. Water samples were collected from a pond adjacent to an experimental forest in the Center for Ecological Research, Kyoto University (34° 58′ 18″ N, 135° 57′ 33″ E) in February 2017. Ten ml of water samples were collected from the pond and filtered using the cartridge filters. The amounts of beads inside the filter cartridge were 0 (No beads), 0.05, 0.2, 1 and 2 g. The number of replicates for each category was three. In total, 15 samples were included in the preliminary test (i.e., three replicate and five treatments). After the filtration, DNAs were extracted using a DNeasy® Blood & Tissue Kit (Qiagen, Hilden, Germany) as described in the main text (see DNA extraction in Methods section). The yields of extracted DNA were quantified using a NanoDrop spectrometer (ThermoFisher Scientific, Waltham, Massachusetts, USA).

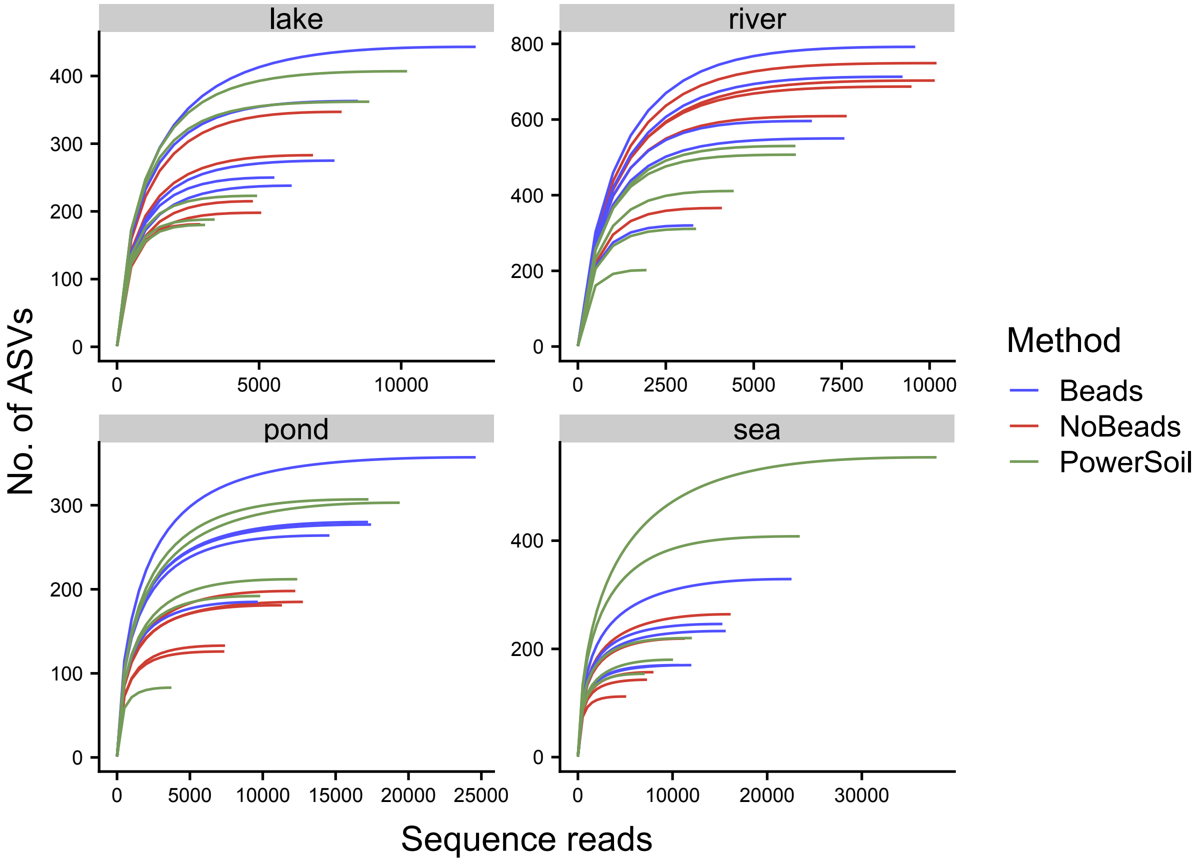


**Figure S2**| Rarefaction curve of sequence reads of each sample. Different colours indicate different DNA extraction methods.

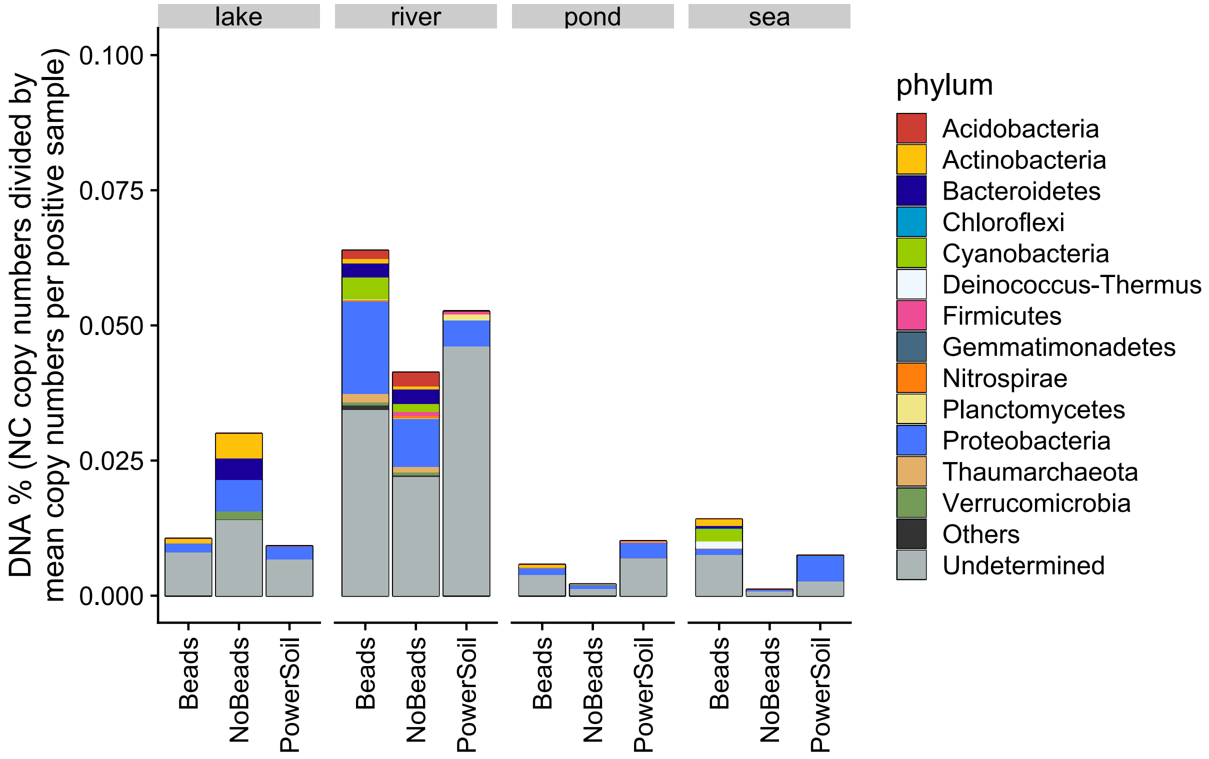


**Figure S3**| Sequence reads detected in the field negative controls. The DNA copy numbers detected in a field negative control were divided by the mean DNA copy number of positive samples in each method. DNA copy numbers detected in field negative controls were less than 0.07%. In addition, there were no qualitative differences in DNA copy numbers or phylum detected among DNA extraction methods.

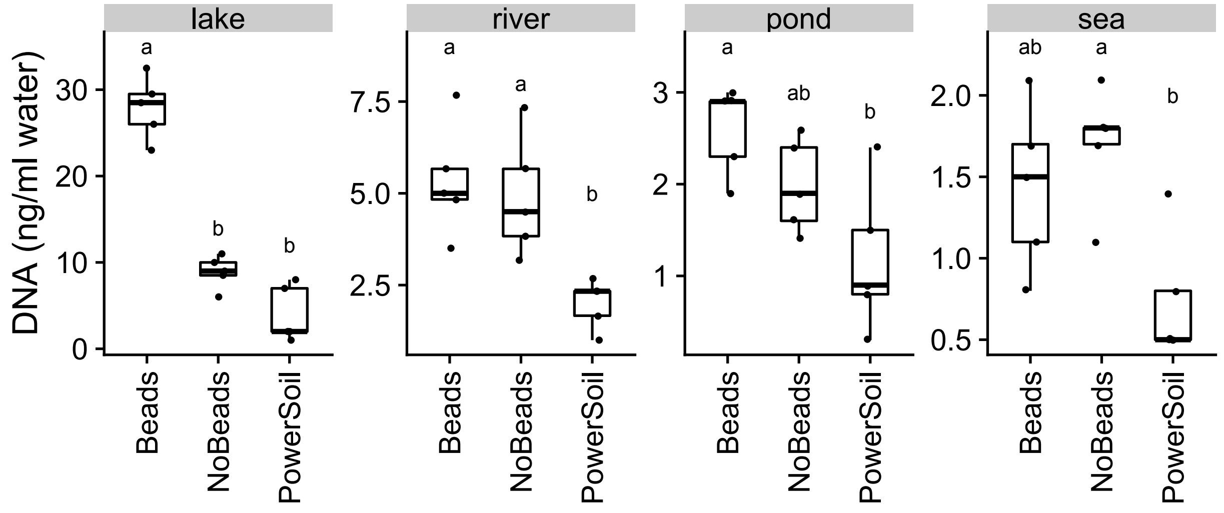


**Figure S4**| The relationship between DNA yield quantified by NanoDrop and DNA extraction method for the lake and river samples. Different letters indicate significant differences between the DNA extraction methods ($P < 0.05$).

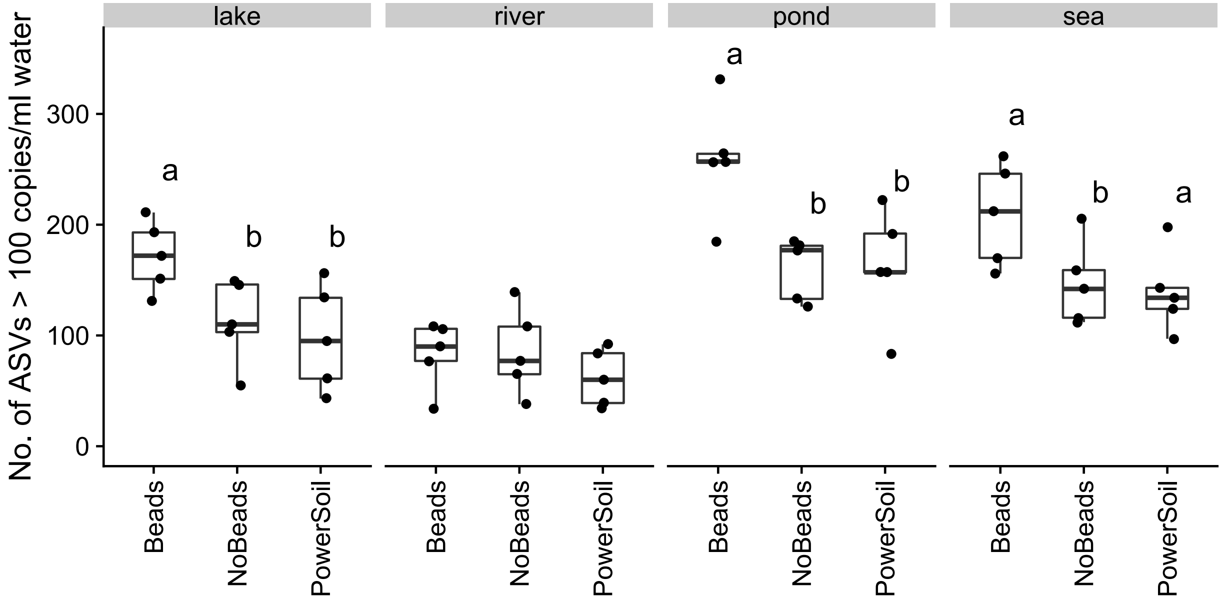


**Figure S5**| The number of ASVs with more than 100 copies per ml water for each treatment. Different letters indicate significant differences between the DNA extraction methods ($P < 0.05$).

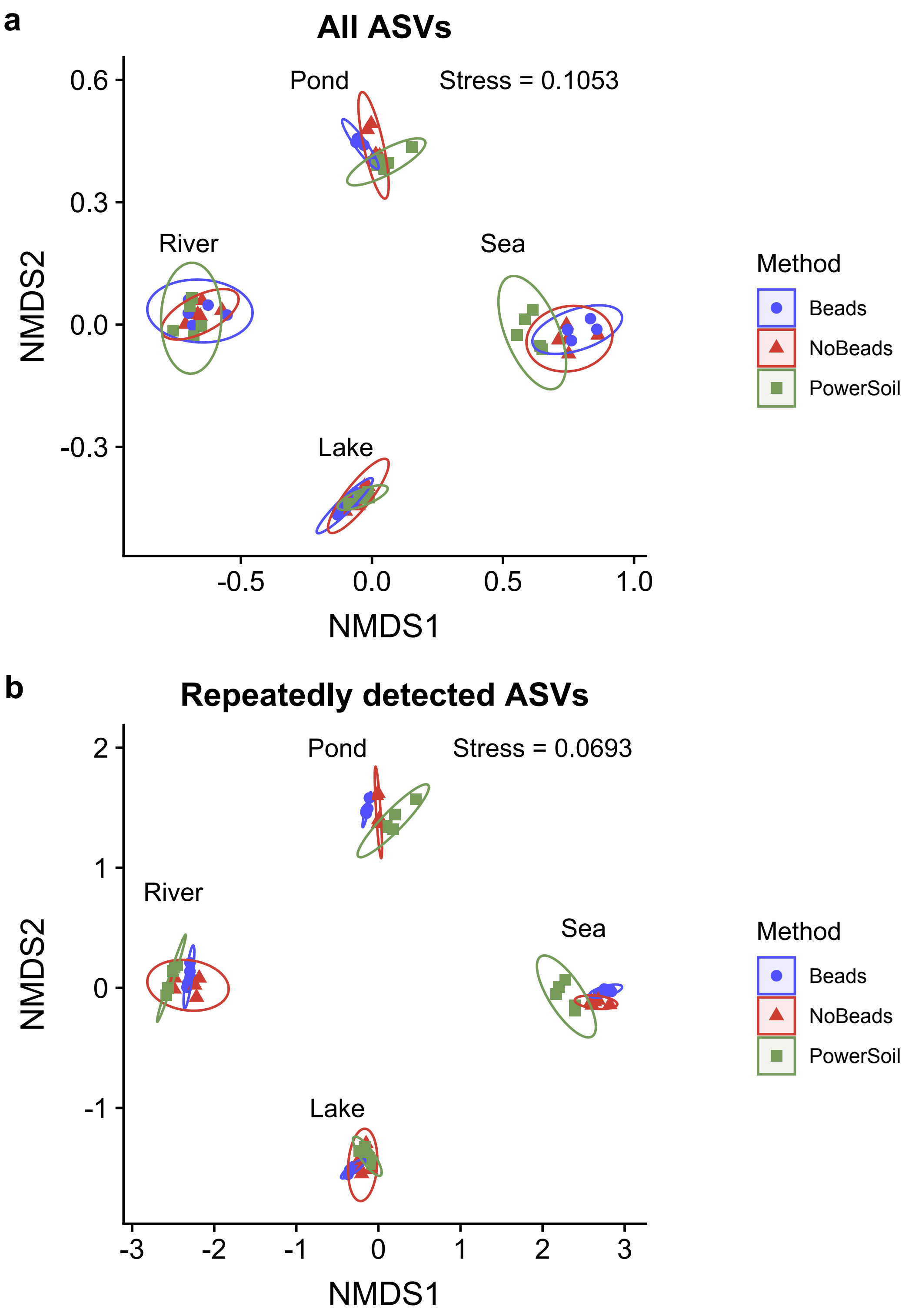


**Figure S6**| Nonmetric dimensional scaling (NMDS) of the prokaryotic community composition. All ASVs from all sites were used for (**a**) and only repeatedly detected ASVs from all sites were used for (**b**). "*P*" indicates the significance of the difference among methods in the overall community composition.

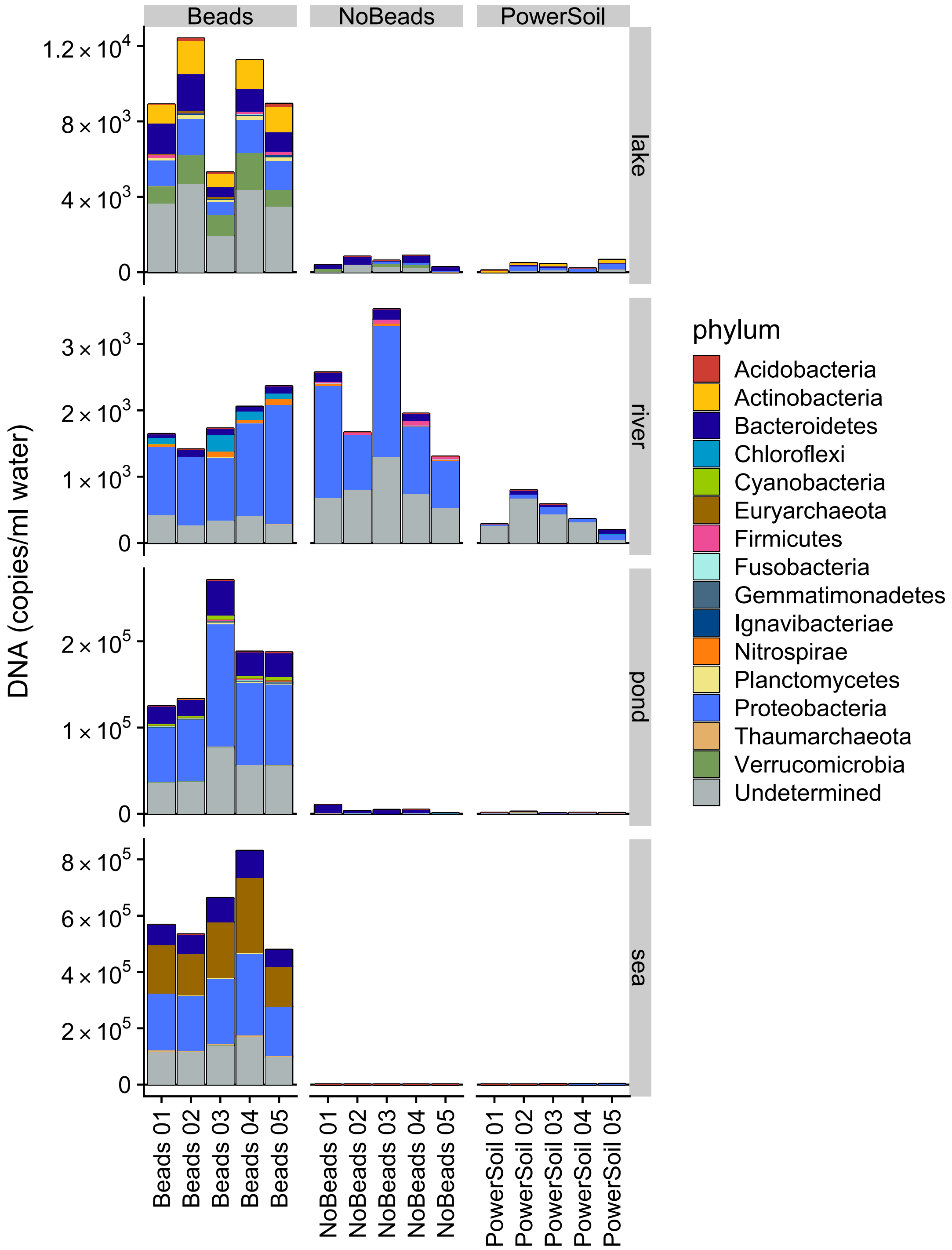


**Figure S7**| Method-specific ASVs for each DNA extraction method based on an alternative, quantitative criterion. The alternative, more quantitative criterion was applied and used to test whether the results remained qualitatively unchanged. First, mean value, standard deviation, and coefficient of variation (C.V.) of each microbial ASV were calculated. Then, the criterion for method-specific ASV was as follows: (1) the mean value of the method-specific ASV is two times as large as that of the ASV detected by other methods and (2) the C.V. of the method-specific ASV is less than one. ASVs satisfying criteria (1) and (2) are ASVs that are predominantly and reliably detected by a method, and thus the combination of these can be an alternative criterion for method-specific ASVs.

**Table S1| Sample metadata and sequence reads passed DADA2 processing**

| *Sample metadata* | | | | | | | | *DADA2 processing* | | | | | | | | |
|---|---|---|---|---|---|---|---|---|---|---|---|---|---|---|---|---|
| **Sample ID** | **Site** | **DNA extraction method** | **Sample or NC** | **Filtered water volume (ml)** | **Sampling date** | **i7 tag sequence (rev. comp.)** | **i5 tag sequence** | **Input (reads)** | **Quality filtered (reads)** | **Denoised (reads)** | **Pair-reads merged (reads)** | **Chimera removed (reads)** | **Input/ Nonchimera (%)** | **Sum of STD seqs (reads)** | **Sum of non-STD seqs (reads)** | **Non STD/total seqs (%)** |
| ***Samples analyzed in the 1st MiSeq run*** | | | | | | | | | | | | | | | | |
| S001 | lake | No Beads | Sample | 20 | 2017/02/21 | GCATGCAT | TATAGCCT | 35,185 | 33,457 | 32,619 | 31,567 | 31,522 | 89.6 | 19,714 | 11,808 | 37.5 |
| S002 | lake | No Beads | Sample | 20 | 2017/02/21 | GCATGCAT | ATAGAGGC | 28,400 | 26,095 | 25,342 | 24,567 | 24,535 | 86.4 | 17,252 | 7,283 | 29.7 |
| S003 | lake | No Beads | Sample | 20 | 2017/02/21 | GCATGCAT | CCTATCCT | 31,330 | 30,074 | 29,310 | 28,418 | 28,393 | 90.6 | 18,779 | 9,614 | 33.9 |
| S004 | lake | No Beads | Sample | 20 | 2017/02/21 | GCATGCAT | GGCTCTGA | 25,709 | 24,792 | 24,139 | 23,472 | 23,453 | 91.2 | 16,263 | 7,190 | 30.7 |
| S005 | lake | No Beads | Sample | 20 | 2017/02/21 | GCATGCAT | AGGCGAAG | 30,543 | 29,648 | 29,277 | 28,777 | 28,735 | 94.1 | 24,356 | 4,379 | 15.2 |
| S006 | lake | No Beads | Field NC | 20 | 2017/02/21 | GCATGCAT | TAATCTTA | 37,389 | 36,595 | 36,418 | 36,328 | 36,221 | 96.9 | 35,536 | 685 | 1.9 |
| S007 | lake | Beads | Sample | 20 | 2017/02/21 | GCATGCAT | CAGGACGT | 24,502 | 23,259 | 22,419 | 21,558 | 21,537 | 87.9 | 12,934 | 8,603 | 39.9 |
| S008 | lake | Beads | Sample | 20 | 2017/02/21 | GCATGCAT | GTACTGAC | 24,130 | 22,166 | 21,130 | 20,265 | 20,256 | 83.9 | 10,424 | 9,832 | 48.5 |
| S009 | lake | Beads | Sample | 20 | 2017/02/21 | GCCGTAAT | TATAGCCT | 44,290 | 42,314 | 41,531 | 40,473 | 40,412 | 91.2 | 26,874 | 13,538 | 33.5 |
| S010 | lake | Beads | Sample | 20 | 2017/02/21 | GCCGTAAT | ATAGAGGC | 29,959 | 27,635 | 26,592 | 25,542 | 25,540 | 85.2 | 13,534 | 12,006 | 47.0 |
| S011 | lake | Beads | Sample | 20 | 2017/02/21 | GCCGTAAT | CCTATCCT | 47,079 | 45,116 | 44,060 | 42,623 | 42,595 | 90.5 | 23,600 | 18,995 | 44.6 |
| S012 | lake | Beads | Field NC | 20 | 2017/02/21 | GCCGTAAT | GGCTCTGA | 28,033 | 25,278 | 25,190 | 25,138 | 25,098 | 89.5 | 25,020 | 78 | 0.3 |
| S013 | lake | PowerSoil | Sample | 20 | 2017/02/21 | GCCGTAAT | AGGCGAAG | 36,265 | 34,954 | 34,571 | 34,064 | 34,036 | 93.9 | 29,497 | 4,539 | 13.3 |
| S014 | lake | PowerSoil | Sample | 20 | 2017/02/21 | GCCGTAAT | TAATCTTA | 48,306 | 46,467 | 45,613 | 44,221 | 44,154 | 91.4 | 28,210 | 15,944 | 36.1 |
| S015 | lake | PowerSoil | Sample | 20 | 2017/02/21 | GCCGTAAT | CAGGACGT | 34,738 | 32,395 | 31,821 | 31,150 | 31,122 | 89.6 | 23,423 | 7,699 | 24.7 |
| S016 | lake | PowerSoil | Sample | 20 | 2017/02/21 | GCCGTAAT | GTACTGAC | 36,328 | 33,918 | 33,545 | 32,862 | 32,737 | 90.1 | 27,709 | 5,028 | 15.4 |
| S017 | lake | PowerSoil | Sample | 20 | 2017/02/21 | CTAGGTTG | TATAGCCT | 49,011 | 46,345 | 45,464 | 44,091 | 44,010 | 89.8 | 30,829 | 13,181 | 30.0 |
| S018 | lake | PowerSoil | Field NC | 20 | 2017/02/21 | CTAGGTTG | ATAGAGGC | 23,515 | 16,676 | 16,532 | 16,503 | 16,490 | 70.1 | 16,456 | 34 | 0.2 |
| S019 | river | No Beads | Sample | 30 | 2017/02/21 | CTAGGTTG | CCTATCCT | 52,250 | 48,773 | 46,643 | 42,883 | 42,798 | 81.9 | 30,169 | 12,629 | 29.5 |
| S020 | river | No Beads | Sample | 30 | 2017/02/21 | CTAGGTTG | GGCTCTGA | 51,647 | 49,484 | 47,890 | 44,857 | 44,765 | 86.7 | 34,994 | 9,771 | 21.8 |
| S021 | river | No Beads | Sample | 30 | 2017/02/21 | CTAGGTTG | AGGCGAAG | 50,321 | 47,962 | 45,877 | 41,741 | 41,661 | 82.8 | 28,666 | 12,995 | 31.2 |
| S022 | river | No Beads | Sample | 30 | 2017/02/21 | CTAGGTTG | TAATCTTA | 60,220 | 57,501 | 55,710 | 52,017 | 51,958 | 86.3 | 38,706 | 13,252 | 25.5 |
| S023 | river | No Beads | Sample | 30 | 2017/02/21 | CTAGGTTG | CAGGACGT | 44,144 | 42,379 | 41,241 | 39,462 | 39,380 | 89.2 | 34,033 | 5,347 | 13.6 |
| S024 | river | No Beads | Field NC | 20 | 2017/02/21 | CTAGGTTG | GTACTGAC | 47,423 | 45,595 | 45,202 | 45,091 | 44,929 | 94.7 | 44,598 | 331 | 0.7 |
| S025 | river | Beads | Sample | 30 | 2017/02/21 | GACTTGTG | TATAGCCT | 44,754 | 42,192 | 40,528 | 38,176 | 38,027 | 85.0 | 29,155 | 8,872 | 23.3 |
| S026 | river | Beads | Sample | 30 | 2017/02/21 | GACTTGTG | ATAGAGGC | 41,993 | 37,584 | 36,131 | 34,135 | 34,056 | 81.1 | 29,473 | 4,583 | 13.5 |
| S027 | river | Beads | Sample | 30 | 2017/02/21 | GACTTGTG | CCTATCCT | 50,940 | 48,379 | 46,448 | 43,250 | 43,144 | 84.7 | 30,023 | 13,121 | 30.4 |
| S028 | river | Beads | Sample | 30 | 2017/02/21 | GACTTGTG | GGCTCTGA | 50,122 | 47,477 | 45,160 | 41,323 | 41,249 | 82.3 | 29,024 | 12,225 | 29.6 |
| S029 | river | Beads | Sample | 30 | 2017/02/21 | GACTTGTG | AGGCGAAG | 48,030 | 44,375 | 42,598 | 38,740 | 38,703 | 80.6 | 28,882 | 9,821 | 25.4 |
| S030 | river | Beads | Field NC | 20 | 2017/02/21 | GACTTGTG | TAATCTTA | 55,340 | 54,036 | 53,684 | 53,619 | 53,360 | 96.4 | 52,713 | 647 | 1.2 |
| S031 | river | PowerSoil | Sample | 30 | 2017/02/21 | GACTTGTG | CAGGACGT | 40,793 | 38,535 | 37,356 | 35,760 | 35,686 | 87.5 | 31,332 | 4,354 | 12.2 |
| S032 | river | PowerSoil | Sample | 30 | 2017/02/21 | GACTTGTG | GTACTGAC | 42,681 | 37,488 | 35,588 | 33,029 | 32,948 | 77.2 | 24,980 | 7,968 | 24.2 |
| S033 | river | PowerSoil | Sample | 30 | 2017/02/21 | CACTAGTG | TATAGCCT | 38,246 | 36,223 | 34,601 | 32,359 | 32,323 | 84.5 | 24,676 | 7,647 | 23.7 |
| S034 | river | PowerSoil | Sample | 30 | 2017/02/21 | CACTAGTG | ATAGAGGC | 27,472 | 25,423 | 24,326 | 23,146 | 23,114 | 84.1 | 20,582 | 2,532 | 11.0 |
| S035 | river | PowerSoil | Sample | 30 | 2017/02/21 | CACTAGTG | CCTATCCT | 36,742 | 35,475 | 34,124 | 32,251 | 32,165 | 87.5 | 26,598 | 5,567 | 17.3 |
| S036 | river | PowerSoil | Field NC | 20 | 2017/02/21 | CACTAGTG | GGCTCTGA | 38,344 | 37,558 | 37,548 | 37,520 | 37,437 | 97.6 | 37,209 | 228 | 0.6 |
| S037 | NA | NA | R NC w/ S | NA | NA | CACTAGTG | AGGCGAAG | 40,855 | 40,046 | 40,015 | 39,894 | 39,817 | 97.5 | 39,627 | 190 | 0.5 |
| S038 | NA | NA | R NC w/ S | NA | NA | CACTAGTG | TAATCTTA | 39,040 | 38,236 | 38,217 | 38,169 | 38,022 | 97.4 | 37,805 | 217 | 0.6 |
| S039 | NA | NA | PCR NC | NA | NA | CACTAGTG | CAGGACGT | 1,454 | 1,219 | 1,214 | 1,214 | 1,214 | 83.5 | 3 | 1,211 | 99.8 |
| S040 | NA | NA | PCR NC | NA | NA | CACTAGTG | GTACTGAC | 1,418 | 1,138 | 1,131 | 1,110 | 1,110 | 78.3 | 34 | 1,076 | 96.9 |

**Table S1| Continued.**

| *Sample metadata* | | | | | | | | *DADA2 processing* | | | | | | | | |
|---|---|---|---|---|---|---|---|---|---|---|---|---|---|---|---|---|
| **Sample ID** | **Site** | **DNA extraction method** | **Sample or NC** | **Filtered water volume (ml)** | **Sampling date** | **i7 tag sequence (rev. comp.)** | **i5 tag sequence** | **Input (reads)** | **Quality filtered (reads)** | **Denoised (reads)** | **Pair-reads merged (reads)** | **Chimera removed (reads)** | **Input /Nonchimera (%)** | **Sum of STD seqs (reads)** | **Sum of non-STD seqs (reads)** | **Non STD/total seqs (%)** |
| ***Samples analyzed in the 2nd MiSeq run*** | | | | | | | | | | | | | | | | |
| R001 | sea | No Beads | Sample | 100 | 2018/12/27 | GAGACACA | CAGGACGT | 14,602 | 14,309 | 13,547 | 12,966 | 12,932 | 88.6 | 1,401 | 11,531 | 89.2 |
| R002 | sea | No Beads | Sample | 100 | 2018/12/27 | GAGACACA | GTACTGAC | 9,147 | 8,788 | 8,319 | 7,978 | 7,959 | 87.0 | 1,104 | 6,855 | 86.1 |
| R003 | sea | No Beads | Sample | 100 | 2018/12/27 | AACCGGAA | TATAGCCT | 20,593 | 20,175 | 19,220 | 18,600 | 18,518 | 89.9 | 2,701 | 15,817 | 85.4 |
| R004 | sea | No Beads | Sample | 100 | 2018/12/27 | AACCGGAA | ATAGAGGC | 14,628 | 13,815 | 13,009 | 12,496 | 12,479 | 85.3 | 2,759 | 9,720 | 77.9 |
| R005 | sea | No Beads | Sample | 100 | 2018/12/27 | AACCGGAA | CCTATCCT | 27,637 | 27,099 | 25,918 | 24,812 | 24,695 | 89.4 | 2,397 | 22,298 | 90.3 |
| R006 | sea | No Beads | Field NC | 100 | 2018/12/27 | AACCGGAA | GGCTCTGA | 41,080 | 40,322 | 40,296 | 40,216 | 40,029 | 97.4 | 39,352 | 677 | 1.7 |
| R007 | sea | Beads | Sample | 100 | 2018/12/27 | CGAACGAA | CCTATCCT | 25,649 | 25,165 | 23,965 | 22,987 | 22,836 | 89.0 | 1,527 | 21,309 | 93.3 |
| R008 | sea | Beads | Sample | 100 | 2018/12/27 | CGAACGAA | GGCTCTGA | 36,551 | 35,807 | 34,244 | 32,869 | 32,649 | 89.3 | 2,250 | 30,399 | 93.1 |
| R009 | sea | Beads | Sample | 100 | 2018/12/27 | CGAACGAA | AGGCGAAG | 24,279 | 23,806 | 22,708 | 21,885 | 21,741 | 89.5 | 1,244 | 20,497 | 94.3 |
| R010 | sea | Beads | Sample | 100 | 2018/12/27 | CGAACGAA | TAATCTTA | 18,738 | 18,386 | 17,479 | 16,811 | 16,762 | 89.5 | 710 | 16,052 | 95.8 |
| R011 | sea | Beads | Sample | 100 | 2018/12/27 | CGAACGAA | CAGGACGT | 19,614 | 19,138 | 18,098 | 17,174 | 17,088 | 87.1 | 1,436 | 15,652 | 91.6 |
| R012 | sea | Beads | Field NC | 100 | 2018/12/27 | CGAACGAA | GTACTGAC | 33,925 | 31,495 | 31,376 | 31,178 | 30,967 | 91.3 | 27,625 | 3,342 | 10.8 |
| R013 | sea | PowerSoil | Sample | 100 | 2018/12/27 | CTTGGTTC | CAGGACGT | 17,355 | 16,947 | 16,275 | 15,638 | 15,621 | 90.0 | 3,730 | 11,891 | 76.1 |
| R014 | sea | PowerSoil | Sample | 100 | 2018/12/27 | CTTGGTTC | GTACTGAC | 13,624 | 13,061 | 12,464 | 12,103 | 12,090 | 88.7 | 3,920 | 8,170 | 67.6 |
| R015 | sea | PowerSoil | Sample | 100 | 2018/12/27 | CATCGTTC | TATAGCCT | 45,530 | 44,311 | 43,470 | 42,176 | 42,049 | 92.4 | 13,364 | 28,685 | 68.2 |
| R016 | sea | PowerSoil | Sample | 100 | 2018/12/27 | CATCGTTC | ATAGAGGC | 22,376 | 20,378 | 19,588 | 18,940 | 18,924 | 84.6 | 4,830 | 14,094 | 74.5 |
| R017 | sea | PowerSoil | Sample | 100 | 2018/12/27 | CATCGTTC | CCTATCCT | 60,411 | 59,417 | 58,099 | 56,438 | 56,033 | 92.8 | 11,058 | 44,975 | 80.3 |
| R018 | sea | PowerSoil | Field NC | 100 | 2018/12/27 | CATCGTTC | GGCTCTGA | 52,063 | 51,122 | 50,900 | 50,721 | 50,516 | 97.0 | 48,414 | 2,102 | 4.2 |
| R019 | pond | No Beads | Sample | 100 | 2018/12/27 | GAGACACA | TATAGCCT | 10,469 | 10,247 | 9,645 | 9,240 | 9,240 | 88.3 | 1,077 | 8,163 | 88.3 |
| R020 | pond | No Beads | Sample | 100 | 2018/12/27 | GAGACACA | ATAGAGGC | 10,300 | 9,818 | 9,276 | 8,913 | 8,887 | 86.3 | 884 | 8,003 | 90.1 |
| R021 | pond | No Beads | Sample | 100 | 2018/12/27 | GAGACACA | CCTATCCT | 15,080 | 14,836 | 14,091 | 13,451 | 13,436 | 89.1 | 1,079 | 12,357 | 92.0 |
| R022 | pond | No Beads | Sample | 100 | 2018/12/27 | GAGACACA | GGCTCTGA | 16,741 | 16,401 | 15,612 | 15,068 | 15,031 | 89.8 | 1,239 | 13,792 | 91.8 |
| R023 | pond | No Beads | Sample | 100 | 2018/12/27 | GAGACACA | AGGCGAAG | 16,450 | 16,169 | 15,419 | 14,787 | 14,770 | 89.8 | 1,525 | 13,245 | 89.7 |
| R024 | pond | No Beads | Field NC | 100 | 2018/12/27 | GAGACACA | TAATCTTA | 33,177 | 32,607 | 32,583 | 32,548 | 32,407 | 97.7 | 31,678 | 729 | 2.2 |
| R025 | pond | Beads | Sample | 100 | 2018/12/27 | AACCGGAA | AGGCGAAG | 23,715 | 23,302 | 22,024 | 21,105 | 20,909 | 88.2 | 1,527 | 19,382 | 92.7 |
| R026 | pond | Beads | Sample | 100 | 2018/12/27 | AACCGGAA | TAATCTTA | 23,393 | 22,958 | 21,766 | 20,689 | 20,621 | 88.2 | 1,435 | 19,186 | 93.0 |
| R027 | pond | Beads | Sample | 100 | 2018/12/27 | AACCGGAA | CAGGACGT | 19,472 | 19,094 | 17,968 | 17,086 | 16,888 | 86.7 | 653 | 16,235 | 96.1 |
| R028 | pond | Beads | Sample | 100 | 2018/12/27 | AACCGGAA | GTACTGAC | 13,727 | 12,912 | 11,981 | 11,288 | 11,273 | 82.1 | 624 | 10,649 | 94.5 |
| R029 | pond | Beads | Sample | 100 | 2018/12/27 | CGAACGAA | TATAGCCT | 33,664 | 32,687 | 31,043 | 29,585 | 29,255 | 86.9 | 1,724 | 27,531 | 94.1 |
| R030 | pond | Beads | Field NC | 100 | 2018/12/27 | CGAACGAA | ATAGAGGC | 30,550 | 27,202 | 27,154 | 27,129 | 27,013 | 88.4 | 26,328 | 685 | 2.5 |
| R031 | pond | PowerSoil | Sample | 100 | 2018/12/27 | CTTGGTTC | TATAGCCT | 15,091 | 14,699 | 13,801 | 13,106 | 13,045 | 86.4 | 1,280 | 11,765 | 90.2 |
| R032 | pond | PowerSoil | Sample | 100 | 2018/12/27 | CTTGGTTC | ATAGAGGC | 6,201 | 5,265 | 4,665 | 4,470 | 4,470 | 72.1 | 508 | 3,962 | 88.6 |
| R033 | pond | PowerSoil | Sample | 100 | 2018/12/27 | CTTGGTTC | CCTATCCT | 26,723 | 26,278 | 25,466 | 24,597 | 24,512 | 91.7 | 5,627 | 18,885 | 77.0 |
| R034 | pond | PowerSoil | Sample | 100 | 2018/12/27 | CTTGGTTC | GGCTCTGA | 27,989 | 27,583 | 26,491 | 25,491 | 25,269 | 90.3 | 3,473 | 21,796 | 86.3 |
| R035 | pond | PowerSoil | Sample | 100 | 2018/12/27 | CTTGGTTC | AGGCGAAG | 18,190 | 17,925 | 17,206 | 16,544 | 16,509 | 90.8 | 2,970 | 13,539 | 82.0 |
| R036 | pond | PowerSoil | Field NC | 100 | 2018/12/27 | CTTGGTTC | TAATCTTA | 47,052 | 46,279 | 46,252 | 46,181 | 45,857 | 97.5 | 44,739 | 1,118 | 2.4 |
| R037 | NA | NA | R NC w/ S | NA | NA | CATCGTTC | AGGCGAAG | 50,614 | 49,781 | 49,629 | 49,508 | 49,312 | 97.4 | 47,453 | 1,859 | 3.8 |
| R038 | NA | NA | R NC w/ S | NA | NA | CATCGTTC | TAATCTTA | 33,462 | 32,888 | 32,875 | 32,844 | 32,806 | 98.0 | 32,144 | 662 | 2.0 |
| R039 | NA | NA | PCR NC | NA | NA | CATCGTTC | CAGGACGT | 7,571 | 7,433 | 7,309 | 7,213 | 7,213 | 95.3 | 226 | 6,987 | 96.9 |
| R040 | NA | NA | PCR NC | NA | NA | CATCGTTC | GTACTGAC | 4,791 | 4,550 | 4,451 | 4,420 | 4,420 | 92.3 | 187 | 4,233 | 95.8 |
| **Total** | | | | | | | | **2,501,165** | **2,388,717** | **2,322,487** | **2,242,616** | **2,235,743** | **89.4** | **1,431,894** | **803,849** | **36.0** |

**Table S2| Prokaryote standard DNA sequences including primer resions**

| DNA ID | Sequence |
|---|---|
| STD_pro1 | GTGCCAGCAGCCGCGGTAAGACGGAGGGGGCTAGCGTTGTTCGGAATTACTGGGCGTAAA<br>GAGAAGGTAGGCGGAAGCTGAAGTCATGTGTGAAAACGCCTGGCTTAACTTAGCTCAGGG<br>TCGCTAAACTGGTTGGCTTGAGTGTGAACGAGGTCCTCGGAATTTTCTGTGTAGCGGTGA<br>AATGCGTAGATATTAAGGCGAACACCTGCGGCGAAGGCACGGAGCTGGGGCAGGGCTGAC<br>GCTGAGGCGTTAAAGCGTGGGGAGCAAACAGGATTAGATACCCTGGTAGTCC |
| STD_pro2 | GTGCCAGCAGCCGCGGTAAGACGGAGGGGGCTAGCGTTGTTCGGAATTACTGGGCGTAAA<br>GAGTATGTAGGCGGTAAAGGAAGTTACGAGTGAAATTACAGGGCTTAACCGATAAGTCGT<br>GGCCAAAACTGGGAGCCTTGAGTAATCGAGAGGTGGGCGGAATTGGGTGTGTAGCGGTGA<br>AATGCGTAGATATTCAAAGGAACACCGATCGCGAAGGCGGCCTCCTGGTTAGGTCCTGAC<br>GCTGAGGAACGAAAGCGTGGGGAGCAAACAGGATTAGATACCCTGGTAGTCC |
| STD_pro3 | GTGCCAGCAGCCGCGGTAAGACGGAGGGGGCTAGCGTTGTTCGGAATTACTGGGCGTAAA<br>GAGTTAGTAGGCGGGCAATTAAGTTATAGGTGAAAAGTAATGGCTTAACTTCGCGAACTC<br>CGGACAAACTGAGGTGCTTGAGCGTTAAAGAGGCTCCCGGAATTGAAGGTGTAGCGGTGA<br>AATGCGTAGATATGTTTCGGAACACCTAATGCGAAGGCTCAAGTCTGGGTAACTGGTGAC<br>GCTGAGGTCACAAAGCGTGGGGAGCAAACAGGATTAGATACCCTGGTAGTCC |
| STD_pro4 | GTGCCAGCAGCCGCGGTAAGACGGAGGGGGCTAGCGTTGTTCGGAATTACTGGGCGTAAA<br>GAGAGTGTAGGCGGCCACGTAAGTTCCCGGTGAAATCGAGCGGCTTAACGTGCTCCTCGC<br>CCGAGAAACTGAAGCCCTTGAGCTCAGCCGAGGAACCCGGAATTACTTGTGTAGCGGTGA<br>AATGCGTAGATATGTGTATGAACACCTCCAGCGAAGGCCGCACACTGGCACTCCACTGAC<br>GCTGAGGTTTAAAAGCGTGGGGAGCAAACAGGATTAGATACCCTGGTAGTCC |
| STD_pro5 | GTGCCAGCAGCCGCGGTAAGACGGAGGGGGCTAGCGTTGTTCGGAATTACTGGGCGTAAA<br>GAGCCGGTAGGCGGCTCCCGAAGTCGGTAGTGAAATTCTGAGGCTTAACTAAAACCACGA<br>CATTGAAACTGGGTGTCTTGAGTTGATACGAGGCCAGTGGAATTGTGCGTGTAGCGGTGA<br>AATGCGTAGATATAATAAGGAACACCTCCCGCGAAGGCTTTGGCCTGGCATGCCAGTGAC<br>GCTGAGGTGTTAAAGCGTGGGGAGCAAACAGGATTAGATACCCTGGTAGTCC |

**Table S3| Method-specific microbial ASVs**

| Taxa ID | Superkingdom | Phylum | Class | order | family | genus |
|---|---|---|---|---|---|---|
| ***Lake - Beads*** | | | | | | |
| Taxa_00190 | Bacteria | | | | | |
| Taxa_00433 | Bacteria | | | | | |
| Taxa_00792 | Bacteria | Proteobacteria | Deltaproteobacteria | | | |
| Taxa_00797 | Bacteria | Verrucomicrobia | Verrucomicrobiae | Verrucomicrobiales | | |
| Taxa_00886 | Bacteria | Proteobacteria | Gammaproteobacteria | | | |
| Taxa_00893 | Bacteria | Proteobacteria | Alphaproteobacteria | Sphingomonadales | Sphingomonadaceae | |
| Taxa_00917 | Bacteria | Proteobacteria | Alphaproteobacteria | Sphingomonadales | Sphingomonadaceae | |
| Taxa_01004 | Bacteria | Proteobacteria | Betaproteobacteria | | | |
| Taxa_01142 | Bacteria | Ignavibacteriae | Ignavibacteria | Ignavibacteriales | Ignavibacteriaceae | *Ignavibacterium* |
| Taxa_01154 | Bacteria | Proteobacteria | Deltaproteobacteria | | | |
| Taxa_01165 | Bacteria | Proteobacteria | Betaproteobacteria | | | |
| Taxa_01283 | Bacteria | Bacteroidetes | | | | |
| Taxa_01686 | Bacteria | Bacteroidetes | Cytophagia | Cytophagales | Cytophagaceae | *Ohtaekwangia* |
| ***Lake - No Beads*** | | | | | | |
| Taxa_00612 | Bacteria | Bacteroidetes | Flavobacteriia | Flavobacteriales | Flavobacteriaceae | *Flavobacterium* |
| Taxa_00892 | Bacteria | Planctomycetes | Phycisphaerae | Phycisphaerales | Phycisphaeraceae | *Phycisphaera* |
| Taxa_00960 | Bacteria | Bacteroidetes | Cytophagia | Cytophagales | Cytophagaceae | |
| Taxa_00977 | Bacteria | Bacteroidetes | | | | |
| Taxa_01096 | Bacteria | Proteobacteria | Alphaproteobacteria | | | |
| ***Lake - PowerSoil*** | | | | | | |
| Taxa_00327 | Bacteria | | | | | |
| Taxa_00528 | Bacteria | Proteobacteria | Oligoflexia | Bdellovibrionales | Bdellovibrionaceae | *Bdellovibrio* |
| Taxa_00785 | Bacteria | Actinobacteria | Acidimicrobiia | Acidimicrobiales | Acidimicrobiaceae | *Ilumatobacter* |
| ***River - Beads*** | | | | | | |
| Taxa_00202 | Bacteria | Proteobacteria | Betaproteobacteria | Burkholderiales | | |
| Taxa_00390 | Bacteria | Proteobacteria | Gammaproteobacteria | Pseudomonadales | Pseudomonadaceae | *Pseudomonas* |
| Taxa_00500 | Bacteria | Chloroflexi | | | | |
| Taxa_00536 | Bacteria | Proteobacteria | Alphaproteobacteria | Sphingomonadales | Sphingomonadaceae | |
| Taxa_00566 | Bacteria | Proteobacteria | Alphaproteobacteria | Sphingomonadales | Sphingomonadaceae | |
| Taxa_00622 | Bacteria | | | | | |
| Taxa_00627 | Archaea | Crenarchaeota | Thermoprotei | Fervidicoccales | Fervidicoccaceae | *Fervidicoccus* |
| Taxa_00639 | Bacteria | Proteobacteria | Alphaproteobacteria | | | |
| Taxa_00672 | Bacteria | Bacteroidetes | Flavobacteriia | Flavobacteriales | Flavobacteriaceae | *Flavobacterium* |
| Taxa_00811 | Bacteria | | | | | |
| Taxa_00864 | Bacteria | Nitrospirae | Nitrospira | Nitrospirales | Nitrospiraceae | *Nitrospira* |
| Taxa_01832 | Bacteria | | | | | |
| Taxa_03037 | Bacteria | | | | | |
| ***River- No Beads*** | | | | | | |
| Taxa_00109 | Bacteria | Proteobacteria | Betaproteobacteria | Burkholderiales | | |
| Taxa_00363 | Bacteria | | | | | |
| Taxa_00408 | Bacteria | Bacteroidetes | | | | |
| Taxa_00540 | Bacteria | | | | | |
| Taxa_00557 | Bacteria | Proteobacteria | | | | |
| Taxa_00708 | Bacteria | Proteobacteria | Deltaproteobacteria | Myxococcales | | |
| Taxa_00712 | Bacteria | Proteobacteria | Oligoflexia | Bdellovibrionales | Bdellovibrionaceae | *Bdellovibrio* |
| Taxa_00822 | Bacteria | | | | | |
| Taxa_00856 | Bacteria | Firmicutes | Bacilli | Lactobacillales | Lactobacillaceae | *Lactobacillus* |
| Taxa_00919 | Bacteria | | | | | |
| Taxa_00966 | Bacteria | Proteobacteria | | | | |
| Taxa_01066 | Bacteria | | | | | |
| Taxa_01954 | Bacteria | Nitrospirae | Nitrospira | Nitrospirales | Nitrospiraceae | |
| ***River- PowerSoil*** | | | | | | |
| Taxa_01184 | Bacteria | Proteobacteria | | | | |
| ***Pond - Beads*** | | | | | | |
| Taxa_00576 | Bacteria | Proteobacteria | Epsilonproteobacteria | Campylobacterales | Helicobacteraceae | *Sulfuricurvum* |
| Taxa_00584 | Bacteria | | | | | |
| Taxa_00610 | Bacteria | Bacteroidetes | Bacteroidia | Marinilabiliales | | |
| Taxa_00690 | Bacteria | Proteobacteria | Epsilonproteobacteria | Campylobacterales | Campylobacteraceae | *Arcobacter* |
| Taxa_00705 | Bacteria | Proteobacteria | Gammaproteobacteria | | | |
| Taxa_00737 | Bacteria | | | | | |
| Taxa_00782 | Bacteria | Proteobacteria | Betaproteobacteria | Burkholderiales | | |
| Taxa_00880 | Bacteria | Proteobacteria | Deltaproteobacteria | Syntrophobacterales | Syntrophaceae | *Syntrophus* |
| Taxa_00912 | Bacteria | Fusobacteria | Fusobacteriia | Fusobacteriales | Fusobacteriaceae | *Cetobacterium* |
| Taxa_00925 | Bacteria | Proteobacteria | Alphaproteobacteria | Caulobacterales | Caulobacteraceae | |
| Taxa_01042 | Bacteria | Proteobacteria | Oligoflexia | Bdellovibrionales | Bdellovibrionaceae | *Bdellovibrio* |
| Taxa_01091 | Bacteria | Verrucomicrobia | Verrucomicrobiae | Verrucomicrobiales | Verrucomicrobiaceae | |
| Taxa_01093 | Bacteria | Proteobacteria | Betaproteobacteria | Rhodocyclales | Azonexaceae | *Dechloromonas* |

| | | | | | | |
|---|---|---|---|---|---|---|
| Taxa_01174 | Bacteria | Proteobacteria | Deltaproteobacteria | Myxococcales | | |
| Taxa_01280 | Archaea | Thaumarchaeota | | | | *Candidatus Nitrosotenuis* |
| Taxa_01595 | Bacteria | Bacteroidetes | Bacteroidia | Bacteroidales | Porphyromonadaceae | *Petrimonas* |
| Taxa_01682 | Bacteria | Proteobacteria | Alphaproteobacteria | Rickettsiales | Anaplasmataceae | *Anaplasma* |
| Taxa_01872 | Bacteria | | | | | |
| Taxa_02091 | Archaea | | | | | |
| Taxa_02188 | Bacteria | | | | | |
| Taxa_02191 | Bacteria | | | | | |
| Taxa_02192 | Archaea | | | | | |
| ***Pond - No Beads*** | | | | | | |
| Taxa_00821 | Bacteria | Proteobacteria | Epsilonproteobacteria | Campylobacterales | Campylobacteraceae | *Arcobacter* |
| ***Pond - PowerSoil*** | | | | | | |
| Taxa_00014 | Bacteria | | | | | |
| Taxa_01141 | Bacteria | Proteobacteria | Gammaproteobacteria | Pseudomonadales | Moraxellaceae | *Acinetobacter* |
| Taxa_01281 | Bacteria | Actinobacteria | Actinobacteria | Micromonosporales | Micromonosporaceae | *Actinoplanes* |
| ***Sea - Beads*** | | | | | | |
| Taxa_00170 | Bacteria | | | | | |
| Taxa_00177 | Archaea | Euryarchaeota | | | | |
| Taxa_00307 | Bacteria | | | | | |
| Taxa_00353 | Bacteria | Proteobacteria | Alphaproteobacteria | Pelagibacterales | Pelagibacteraceae | *Candidatus Pelagibacter* |
| Taxa_00421 | Bacteria | Proteobacteria | Alphaproteobacteria | Rhodobacterales | Rhodobacteraceae | |
| Taxa_00781 | Bacteria | Proteobacteria | Gammaproteobacteria | Oceanospirillales | Oceanospirillaceae | *Marinomonas* |
| Taxa_00866 | Bacteria | Proteobacteria | Gammaproteobacteria | | | |
| Taxa_00884 | Bacteria | Bacteroidetes | Cytophagia | Cytophagales | Flammeovirgaceae | *Flammeovirga* |
| Taxa_00983 | Bacteria | Actinobacteria | Acidimicrobiia | Acidimicrobiales | Acidimicrobiaceae | *Ilumatobacter* |
| Taxa_01087 | Bacteria | Proteobacteria | Gammaproteobacteria | Oceanospirillales | Halomonadaceae | *Candidatus Portiera* |
| ***Sea - No Beads*** | | | | | | |
| Taxa_00384 | Bacteria | Proteobacteria | Gammaproteobacteria | Pseudomonadales | Pseudomonadaceae | *Pseudomonas* |
| ***Sea - PowerSoil*** | | | | | | |
| Taxa_00115 | Bacteria | Proteobacteria | Alphaproteobacteria | Rhizobiales | Methylobacteriaceae | *Methylobacterium* |
| Taxa_00135 | Bacteria | | | | | |
| Taxa_00396 | Bacteria | Proteobacteria | Gammaproteobacteria | | | |
| Taxa_00751 | Bacteria | Proteobacteria | Alphaproteobacteria | Rhodobacterales | Rhodobacteraceae | |
| Taxa_00759 | Bacteria | Proteobacteria | Gammaproteobacteria | Cellvibrionales | Halieaceae | |
| Taxa_00790 | Bacteria | Actinobacteria | Acidimicrobiia | Acidimicrobiales | Acidimicrobiaceae | *Ilumatobacter* |
| Taxa_01192 | Bacteria | Firmicutes | Clostridia | Clostridiales | Clostridiaceae | *Clostridium* |
| Taxa_02725 | Bacteria | | | | | |
